# Osteoblast-intrinsic defect in glucose metabolism impairs bone formation in type II diabetic mice

**DOI:** 10.1101/2023.01.16.524248

**Authors:** Fangfang Song, Won Dong Lee, Tyler Marmo, Xing Ji, Chao Song, Xueyang Liao, Rebbeca Seeley, Lutian Yao, Haoran Liu, Fanxin Long

**Affiliations:** Translational Research Program in Pediatric Orthopedics, Department of Surgery, The Children’s Hospital of Philadelphia; The State Key Laboratory Breeding Base of Basic Science of Stomatology and Key Laboratory for Oral Biomedicine of Ministry of Education, School and Hospital of Stomatology, Wuhan University, Wuhan, China; Lewis Sigler Institute for Integrative Genomics, Princeton University, Princeton, NJ, USA; Deaprtment of Orthopedic Surgery, University of Pennsylvania, Philadelphia, PA; Department of Computer Science, New Jersey Institute of Technology, Newark, NJ, USA

## Abstract

Skeletal fragility is associated with type 2 diabetes mellitus (T2D), but the underlying mechanism is not well understood. Here, in a mouse model for youth-onset T2D, we show that both trabecular and cortical bone mass are reduced due to diminished osteoblast activity. Stable isotope tracing in vivo with ^13^C-glucose demonstrates that both glycolysis and glucose fueling of the TCA cycle are impaired in diabetic bones. Similarly, Seahorse assays show suppression of both glycolysis and oxidative phosphorylation by diabetes in bone marrow mesenchymal cells as a whole, whereas single-cell RNA sequencing reveals distinct modes of metabolic dysregulation among the subpopulations. Metformin not only promotes glycolysis and osteoblast differentiation in vitro, but also improves bone mass in diabetic mice. Finally, targeted overexpression of Hif1a or Pfkfb3 in osteoblasts of T2D mice averts bone loss. The study identifies osteoblast-intrinsic defects in glucose metabolism as an underlying cause of diabetic osteopenia, which may be targeted therapeutically.

## Introduction

Bone quality is maintained by the coordinated functions of osteoblasts and osteoclasts. During bone formation, polarized osteoblasts secrete extracellular matrix (ECM), consisting of type I collagen and non-collagenous matrix proteins, that mineralizes over time^1, 2^. The production of ECM by osteoblasts is an energy-demanding process and likely requires metabolic reprogramming, but the metabolic regulation of osteoblast differentiation and function is largely unknown^3–5^. Interestingly, prominent signaling pathways that promote or suppress osteogenesis alter glucose metabolism. For instance, parathyroid hormone (PTH), Wnt, insulin-like growth factors (Igf) and Hif1a stimulate glycolysis while promoting osteogenesis, whereas Notch signaling suppresses glycolysis and osteoblast differentiation in mesenchymal progenitors^6–10^. In addition, a number of studies have identified glucose as the main energy substrate for osteoblasts^11–14^. Thus, accumulating evidence to date supports an important role of glucose metabolism in the regulation of osteoblast development and function.

Type 2 diabetes (T2D) is characterized by insulin insensitivity in target tissues resulting in hyperglycemia. Adults with T2D have increased fracture risk despite having normal or increased areal bone mineral density (aBMD)^15–17^. The poor bone quality has been often attributed to abnormal accumulation of advanced glycation end products in the bone matrix, but could also result from reduced bone turnover as indicated by suppression of both bone resorption and formation in T2D^18–20^. Youth-onset T2D has emerged as a significant and increasing health burden in adolescents and young adults^21^. Distinct from adult T2D, T2D in youth has been reported to reduce bone mass due to impaired bone anabolism^22^. Overall, both types of T2D diminish bone formation, but the underlying mechanisms are not fully understood.

Here, we have established and interrogated a mouse model for youth-onset T2D. We show that suppression of glucose metabolism in osteoblasts is the main cause of impaired bone formation in T2D. Importantly, activation of glycolysis in osteoblasts restores bone formation in T2D mice. Together, we identified osteoblast-intrinsic defects in glucose metabolism as an underlying cause of diabetic osteopenia which may be targeted therapeutically.

## Materials and Methods

### Mice

Animal experiments were conducted with approval from the Institutional Animal Use and Care Committee at Children’s Hospital of Philadelphia. Mice were housed with a 12:12h light-dark cycle at 22 °C. The TRE-Pfkfb3 mouse was generated by placing a doxycycline-inducible Pfkfb3 (CDS isoform 6, NCBI NM_133232.3) expression cassette into the Rosa26 locus by homologous recombination. Chimeric founder mice were created by blastocyst injection of the correctly targeted ES cells and then mated with C57BL/6N to confirm germ-line transmission. The other mouse strains including TRE-Glut1, Osx-rtTA, Hif1dPA and TetO-Cre were as previously described, and maintained on the C57BL/6J background^23–26^. For Dox-mediated gene induction, Dox was supplied in drinking water at 1 mg/ml (Doxycycline hyclate, D9891, Sigma).

### T2D mouse model

For T2D induction, C57BL/6J male mice were fed ad libitum a high-fat diet (HFD, Research diet D12492, 60 kcal% Fat) starting at 6 weeks of age, injected with low-dose streptozotocin (STZ, 30 mg/kg, Sigma) once daily for three consecutive days (5-hr fasting before each injection) starting at 12 weeks of age, and continued on HFD until harvest at 22 weeks of age. Males were used because they were more prone to develop T2D with the protocol. The control group was C57BL/6J male mice of the same age and fed ad libitum regular chow (CTRL, Rodent Laboratory Chow 5015, Purina, USA). Both groups had free access to water. Random glucose level was measured at two weeks after the first STZ injection, and mice with levels lower than 200 mg/dL were excluded from further study. Metformin hydrochloride (4 mg/mL, Cayman Chemical, No. 13118) and Dox (1mg/mL) were dissolved in drinking water with the water bottle changed twice weekly.

### GTT and ITT

Glucose tolerance test (GTT) and insulin tolerance test (ITT) were carried out with one week interval for the animals to recover from stress. For ITT, mice were injected intraperitoneally human insulin at 0.5 U/kg body weight (Novalin R U-100) after 6 hours of fasting. Blood samples were collected at indicated time (0, 15, 30, 60 and 90 mins after injection) from a cut in the tail for glucose measurements with a glucometer. For GTT, mice were intraperitoneally injected with glucose at 2 g/kg body weight after 6 hours of fasting. Blood glucose levels were measured at 0, 15, 30, 60, 90 and 120 mins after injection. The upper limit of the glucometer was 600 mg/dL; glucose levels were recorded as 600 mg/dL when the reading was out of range.

### Serum biochemical assays

Sera were collected from mice through retro-orbital bleeding after 6 hours of fasting. CTX-I and P1NP assays were performed with the RatLaps™ (CTX-I) EIA and Rat/Mouse P1NP EIA Kit (Immunodiagnostic Systems, Ltd.), respectively. Insulin and Igf1 assays were performed with mouse insulin ELISA kit (Ultra Sensitive Mouse Insulin ELISA, Cat#90080, Crystal Chem) and mouse Igf1 ELISA kit (Mouse IGF-1 ELISA Kit, Cat#80574, Crystal Chem). HOMA-IR was calculated as follows: HOMA IR = fasting insulin (mU/L) * fasting glucose (mg/dL)/405. Fasting glucose levels were measured with a glucometer in blood collected from a tail-vein cut.

### Skeletal phenotyping

The distal end of the femur was scanned by μCT (μCT45; SCANCO Medical) at 4.5 μm isotropic voxel size. For trabecular bone analyses, regions of interest were selected at 0.45–2.25 mm below the growth plate. For cortical bone, a total of 70 slices at the femur midshaft located at 5.4 mm away from the distal growth plate were analyzed. Parameters computed from these data included bone volume (BV), total volume (TV), BV/TV, trabecular bone thickness (Tb.Th), number (Tb.N), separation (Tb. Sp), and connectivity density (Conn. Dens) at the distal femoral metaphysis and total area (TA), bone area (BA), BA/TA and cortical thickness (Ct.th) at the mid-diaphysis of the femur.

Dynamic histomorphometry was conducted by sequential injections of calcein green (5mg/kg body weight (bw)) and alizarin red (15mg/kg bw) at 7 and 2 days, respectively, prior to sacrifice. Femurs or tibias were fixed with 4% PFA in PBS for 48 hours and then kept in 70% ethanol with protection from light; the fixed bones were switched to 30% sucrose in PBS overnight before OCT embedding and cryostat sectioning. Longitudinal 10 µm cryostat sections of the undecalcified bones were collected with Cryofilm type II membrane (Section Lab, Co. Ltd., Japan) and imaged using Axioscan (ZEISS, Germany). Quantification was performed with BIOQUANT Image Analysis System (BIOQUANT Image Analysis Corporation, USA).

### Stable isotope tracing in vivo

Conscious animals were fasted for 6 hours and then injected with one bolus of ^13^C_6_-Glc (40 mg/mouse) via tail vein at 60 minutes before the plasma and bone shafts were harvested^13^. For plasma metabolite extraction, plasma (5 µl) was added to 120 µl of −20 °C 40:40:20 methanol: acetonitrile: water (extraction solvent), vortexed, and centrifuged at 21,000g for 20 min at 4 °C. The supernatant (40 µl) was collected for liquid chromatography–mass spectrometry (LC–MS) analysis. For bone metabolite extraction, frozen bones (1 tibia and 1 femur) were transferred to 2 ml round-bottom Eppendorf Safe-Lock tubes on dry ice. Samples were then ground into powder with a cryomill machine (Retsch, Germany) for 30 s at 25 Hz, and maintained at a cold temperature using liquid nitrogen. For every 20 mg tissues, 800 μl of −20 °C extraction solvent was added to the tube, vortexed for 10 s, and then centrifuged at 21,000 ×g for 20 min at 4 °C. The supernatants (500 μl) were transferred to another Eppendorf tube, dried using a N2 evaporator (Organomation Associates, Berlin, MA), and redissolved in 50 μl of −20 °C extraction solvent. Redissolved samples were centrifuged at 21,000 ×g for another 20 min at 4 °C. The supernatants were then transferred to plastic vials for LC-MS analysis. A procedure blank was generated identically without tissue, and was used later to remove the background ions.

For metabolite measurements by LC-MS, a quadrupole-orbitrap mass spectrometer (Q Exactive Plus, Thermo Fisher Scientific, San Jose, CA) operating in negative mode was coupled to hydrophobic interaction chromatography (HILIC) via electrospray ionization. Scans were performed from m/z 70 to 1000 at 1 Hz and 140,000 resolution. LC separation was conducted on a XBridge BEH Amide column (2.1 mm x 150 mm x 2.5 mm particle size, 130 Å pore size; Water, Milford, MA) using a gradient of solvent A (20 mM ammonium acetate, 20 mM ammounium hydroxide in 95:5 water: acetonitrile, pH 9.45) and solvent B (acetonitrile). Flow rate was 150 mL/min. The LC gradient was: 0 min, 85% B; 2 min, 85% B; 3 min, 80% B; 5 min, 80% B; 6 min, 75% B; 7 min, 75% B; 8 min, 70% B; 9 min, 70% B; 10 min, 50% B; 12 min, 50% B; 13 min, 25% B; 16 min, 25% B; 18 min, 0% B; 23 min, 0% B; 24 min, 85% B. Autosampler temperature was 5°C, and injection volume was 15 μL. For improved detection of fructose-1,6-bisphosphate and 3-phosphoglycerate, selected ion monitoring (SIM) scans were added. For 3-phosphoglycerate and fructose-1,6-bisphosphate, scans were performed from 180-190 m/z and 336-350 m/z, respectively, from 13-15 min of the 25-min gradient run with 70,000 resolution and maximum IT of 500 ms and AGC target of 3 x 10^6^. Data were analyzed using the El-MAVEN (Elucidata) software ^27^. Carbon Enrichment was calculated as follows: Carbon Enrichment = (m0*0+m1*1+m2*2+….mn*n)/n.

### scRNA-Seq

For isolation of bone marrow mesenchymal cells, epiphyseal ends of both tibias and femurs were removed and the bone marrow cells were flushed out with M-199 (Thermo Fisher Scientific, USA) supplemented with 10% FBS (heat-inactivated Qualified FBS, Thermo Fisher Scientific, USA) and collected. The bone shaft was then cut into pieces and digested with 1 mg/mL STEMxyme1 (Worthington, Ref#LS004106) and 1 mg/mL Dispase II (ThermoFisher Scientific, Ref#17105041), in M-199 supplemented with 2% FBS for 25 min at 37°C. The digested cells and bone marrow cells were pooled together and passed through a 70mm filter. Cells were then stained in media 199 supplemented with 2% FBS for FACS. The antibodies and staining reagents included Ter119-APC (eBioscience, Ref#17-5921-82, clone TER-119), CD71-PECy7 (Biolegend, Ref#113812, clone RI7217), CD45-PE (eBioscience, Ref#12-0451-82, clone 30-F11), CD3-PE (Biolegend, Ref#100206, clone 17A2), B220-PE (Biolegend, Ref#103208, clone RA3-6B2), CD19-PE (Biolegend, Ref#115508, clone 6D5), Gr-1-PE (Biolegend, Ref#108408, clone RB6-8C5), Cd11b-PE (Biolegend, Ref#101208, clone M1/70) and Calcein AM (Thermo Fisher Scientific, Massachusetts, USA) for 30 minutes on ice. 7-AAD (Thermo Fisher Scientific, Massachusetts, USA) was added into the medium before sorting on MoFlo Astrios (Beckman Coulter, Brea, CA, USA). Bone marrow mesenchymal cells were enriched as the 7-AAD^-^ /Calcein AM^+^/ CD71^-^/ Ter119^-^/ Lin (CD45/CD3/B220/CD19/Gr-1/CD11b)^-^ fraction. The cells were pooled from three T2D or CTRL mice. scRNA-seq libraries were constructed with T2D versus CTRL bone marrow mesenchymal cells with the Single Cell 3’ Reagent Kits v3.1 (10x Genomics). Sequencing was performed with a NovaSeq sequencer in a paired-end, single indexing run, generating ∼200 million reads per sample. Cell ranger (10 × Genomics) was used for demultiplexing, extraction of cell barcode and quantification of unique molecular identifiers (UMIs). 8608 and 8629 cells were recovered from CTRL and T2D samples, respectively. Downstream analyses were performed using Seurat v3 ^28^. Cells with poor quality (genes > 6000, genes < 200, or >10% mitochondrial genes) were excluded. We then normalized the data by the total expression, multiplied by a scale factor of 10,000, and log-transformed the result. The medium number of features detected per cell was 2088 and 2333 from CTRL and T2D, respectively. For the integrated datasets, anchors were defined using the FindIntegrationAnchors function, and these anchors from the different datasets were then used to integrate datasets together with IntegrateData. Different resolutions were used to demonstrate the robustness of clusters. In addition, differentially expressed genes within each cluster between two datasets were identified using FindAllMarkers. Gene Set Enrichment Analysis (GSEA) with MsigDB gene sets was performed to identify pathways with significant changes.

### BMSC isolation and purification

Tibias and femurs were dissected clean from adjacent tissues, and the epiphyses containing the growth plates were excised off with scissors. The bone marrow cells were flushed out with a syringe filled with MEM-α media containing 10% FBS and then passed through a cell strainer (35 µm). The cells for all four bones of one mouse were seeded into one 100 mm dish. The culture medium (MEM-α containing 10% FBS) was changed every day after 4 days following seeding, and the cells normally became confluent at day 6-8, at which time they were dissociated using Collagenase II (4 mg/ml) and 0.05 % trypsin with EDTA for 5 mins at 37°C. Hematopoietic lineage cells were immuno-depleted using the MACS technique (Miltenyi Biotec, USA). Briefly, the cells were mixed with CD45 antibody-coated magnetic beads (CD45 MicroBeads, 130-052-301) and then passed through a MACS LD column (Miltenyi Biotec, 130-042-901) on a MidiMACS^TM^ separator (Miltenyi Biotec, 130-042-302) attached to a multistand. The purified cells were cultured and passaged once for osteogenic differentiation and metabolic assays.

For osteoblast differentiation, when the BMSCs reached 100% confluency, MEM-α containing 4 mM ß-glycerol phosphate (Sigma, G9422) and 50 ug/ml ascorbic acid (Sigma, A4544) was added and then changed daily. Cells after 4 or 7 days of differentiation were dissociated with 4 mg/ml collagenase I in PBS for 45 mins, then with 0.05% trypsin for 10 mins for RNA extraction or metabolic assays.

### In vitro metabolic assays

For Seahorse assays, BMSC purified with MACS were seeded at 2 × 10^5^ cells/cm^2^ into XF96 tissue culture microplates (Agilent) at 24 hours prior to experiments. Complete Seahorse medium was prepared from Agilent Seahorse XF Base Medium (Agilent, 102353) containing 5.5 mM glucose, 2 mM glutamine and 1 mM pyruvate, with pH 7.4. The cells were incubated in 180 μl complete Seahorse medium at 37°C for 1 hour in CO_2_-free incubator before measurements in Seahorse XFe96 Analyzer. The following working concentrations of compounds were used: 2 μM Oligomycin, 3 μM FCCP, 1 μM Rotenone, and 1 uM Antimycin A. The oxygen consumption rate (OCR) and extracellular acidification rate (ECAR) were normalized to seeded cell number. Theoretical ATP production rates from glycolysis and OXPHOS were calculated based on the reading of OCR and ECAR. Wave Pro software was used to calculate the following parameters: Proton efflux rate (PER), glycoPER, glycoATP Production Rate, mitoATP Production Rate and TotalATP Production Rate.

For glucose consumption and lactate production, BMSC were seeded at 4 x 10^5^ cells/cm^2^ in 6-well plates and cultured in a customized medium (with glucose, pyruvate and glutamine reconstituted before use) for 24 hours. The medium was collected for glucose or lactate measurement with Glucose (HK) Assay Kit (Sigma, GAHK20) or L-Lactate Assay Kit I (Eton Bioscience, 120001), respectively. The cells were dissociated and counted for normalization.

### RT-qPCR

Total RNA was harvested from cells or bone shaft using RNeasy mini kit (QIAGEN) according to the manufacturer’s protocol. 1 µg RNA was reverse transcribed into cDNA using High-Capacity cDNA Reverse Transcription Kit (Thermo Fisher Scientific, Massachusetts, USA). The gene expression level relative to β-actin was determined by qPCR with gene-specific primers (Supplementary Table S1) and PowerUp SYBR Green Master Mix (Thermo Fisher Scientific, USA). 2^-ΔΔCT^ was calculated for relative fold change.

### Western blot analyses

Protein extracts from bone were prepared in M-PER buffer (cat#78501, Thermo Fisher Scientific, USA) containing phosphatase and proteinase inhibitors (cat#78442, Halt™ Protease and Phosphatase Inhibitor Single-Use Cocktail, Thermo Fisher Scientific, USA). After surgical removal of the epiphysis and flushing of the marrow, the bone shafts of femurs and tibias were snap frozen in liquid nitrogen and then homogenized with a Precellys homogenizer (Bertin Instrument, France). Proteins were separated by electrophoresis in NuPAGE™ 4 to 12% Bis-Tris Mini Protein Gels (Thermo Fisher Scientific, USA), and then transferred to Immobilon®-FL PVDF Membrane (Millipore, USA). The membranes were blocked with Odyssey Blocking Buffer (PBS) (LI-COR Biosciences, USA), and then incubated with primary antibodies overnight. The primary antibodies included Anti-Glucose Transporter Glut1 antibody [EPR3915] (ab115730, 1:2000), Anti-Hexokinase II antibody [EPR20839] (ab209847, 1:1000), Anti-Lactate Dehydrogenase antibody [EP1566Y] (ab52488, 1:1000) and Anti-beta Actin antibody [AC-15] (ab6276, 1:2000). The secondary antibodies IRDye® 800CW Donkey anti-Rabbit IgG (H + L) (LI-COR Biosciences, 1: 20,000) and IRDye® 680RD Goat anti-Mouse IgG (H + L) (LI-COR Biosciences, 1:20,000) were used. The protein bands were imaged with Odyssey® DLx Imaging System and quantified with Image J.

### Statistics

Statistics was conducted with unpaired Student’s t-test, one-way ANOVA followed by Student’s t test between multiple groups or two-way ANOVA with either Sidak’s multiple comparisons or Fisher’s LSD test, as indicated in the figure legends, by using Prism Software 9.0. p value < 0.05 is considered significant. All quantitative data are presented as mean ± SD.

## Results

### Modeling youth-onset T2D in the mouse

To generate a mouse model for human youth-onset T2D, 6-week-old C57BL/6J male mice were fed a high-fat diet (HFD) for 6 weeks and then injected with a low dose of streptozotocin (STZ) once daily for three consecutive days, followed by continuous HFD feeding until harvest at 22 weeks of age (Fig. 1A). Random glucose levels were checked two weeks after the first STZ injection and mice with a glucose level lower than 200 mg/dL were excluded from the study. At harvest, the T2D mice had a significantly higher body weight due to increased fat mass with no change in lean mass (Fig. 1B, C, S1A). They exhibited notable hyperglycemia and significant impairment in glucose handling when compared to the controls, as revealed by glucose tolerance test (GTT) and insulin tolerance test (ITT) (Fig. 1D-I). Fasting serum insulin and Igf1 concentrations were significantly higher in the T2D mice, confirming that they were less sensitive to insulin and reflecting an early phase of T2D (Fig. 1J-K). Their insulin resistance was further demonstrated by the elevated index of Homeostatic Model Assessment for Insulin Resistance (HOMA-IR) (Fig. 1L). Overall, the mouse model exhibits key features of T2D including obesity, hyperglycemia, hyperinsulinemia (early feature) and insulin resistance.

**Figure 1.**
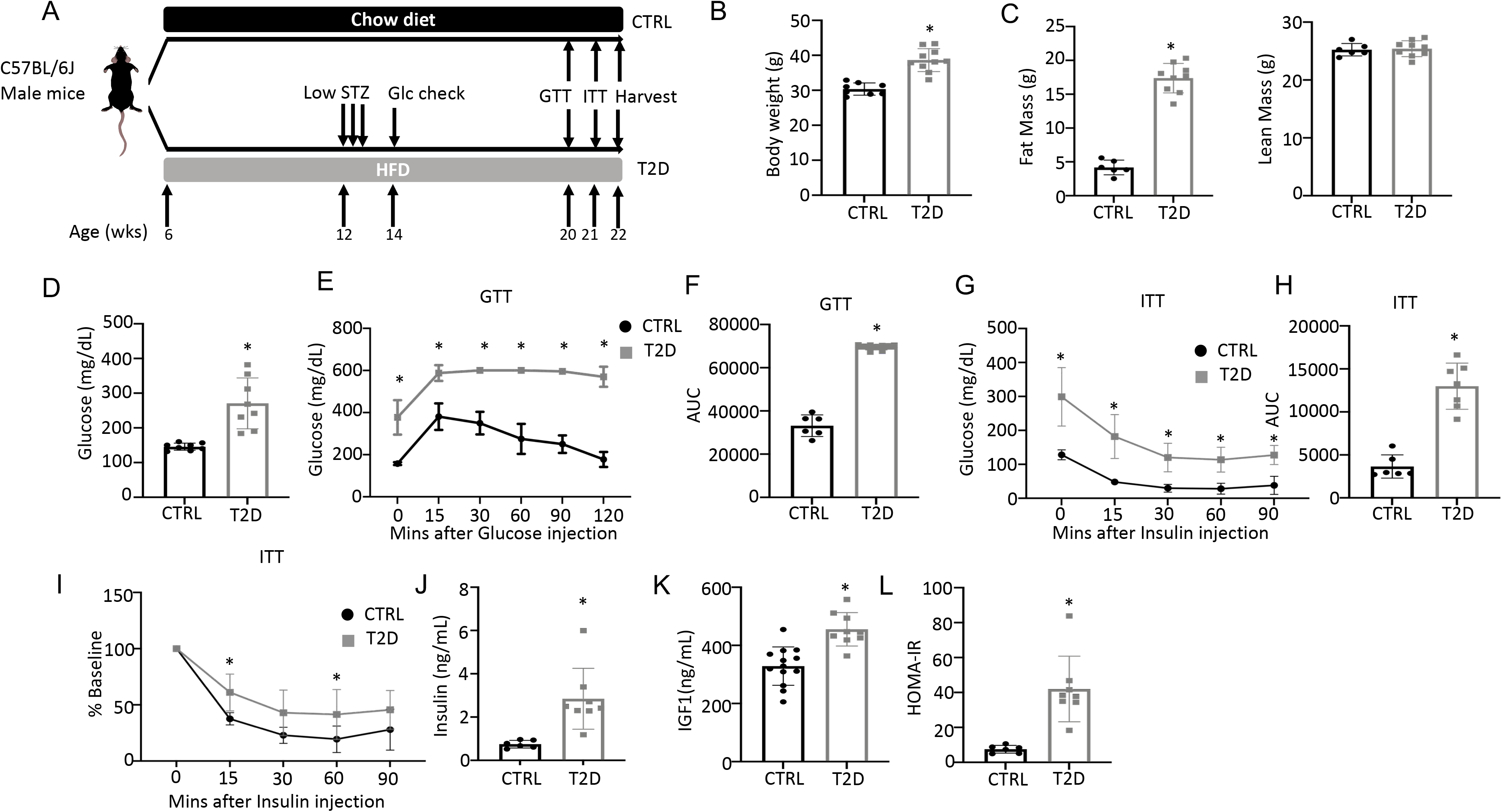
Combination of HFD and low-dose STZ induces T2D in mice. (A) A schematic for experimental design. Low-dose STZ: 30 mg/kg body weight. (B, C) Body weight (B, CTRL, n=8; T2D, n=10) and body composition by DXA (C, CTRL, n=6; T2D, n=9) at harvest. (D) Glucose levels after 6 hr fasting immediately before harvest. n=8. (E, F) Glucose tolerance test (GTT) curve (E) and area under curve (AUC) (F). CTRL, n=6; T2D, n=9. (G, H) Insulin tolerance test (ITT) glucose curve (G), AUC (H) and percentage of baseline (I). CTRL, n=6; T2D, n=7. (J, K) Serum insulin (J, CTRL, n=6; T2D, n=8) and Igf1 (K, CTRL, n=9; T2D, n=13) levels. (L) HOMA-IR graphs. CTRL, n=6; T2D, n=8. All bar graphs are presented as mean ± SD. unpaired student’s t test (B–D, F, H, J-L) or two-way ANOVA with Sidak’s multiple comparisons test (E, G, I). *: P < 0.05.

### T2D reduces bone mass by suppressing osteoblast activity

We next examined bone phenotypes in the T2D mice. Dual-energy x-ray absorptiometry (DEXA) at harvest detected a clear decrease in bone mineral density (BMD) of both whole body (minus the head) and the hindlimb in comparison with the control (Fig. 2A). The body length from nose tip to tail base and the femur length were slightly longer in T2D than normal, perhaps due to increased Igf1 levels (Fig. S1B, C). Imaging and quantification with µCT revealed that T2D significantly reduced trabecular bone volume (BV) and its ratio over tissue volume (BV/TV) in the femur, without altering the tissue volume (TV) itself (Fig. 2B upper, Fig. 2C, Fig. S1D). The trabecular bone loss was due to reduced trabecular number (Tb.N) and connectivity (Conn. Dens) concurrent with increased trabecular spacing (Tb. Sp) (Fig. S1E-G). Similarly, in cortical bone, T2D decreased bone area relative to total area (BA/TA) as well as cortical bone thickness (Ct. Th) at the diaphysis of the femur (Fig. 2B lower, Fig. 2D, Fig. S1H-I). Thus, like human patients, mice with youth-onset T2D exhibit osteopenia.

**Figure 2.**
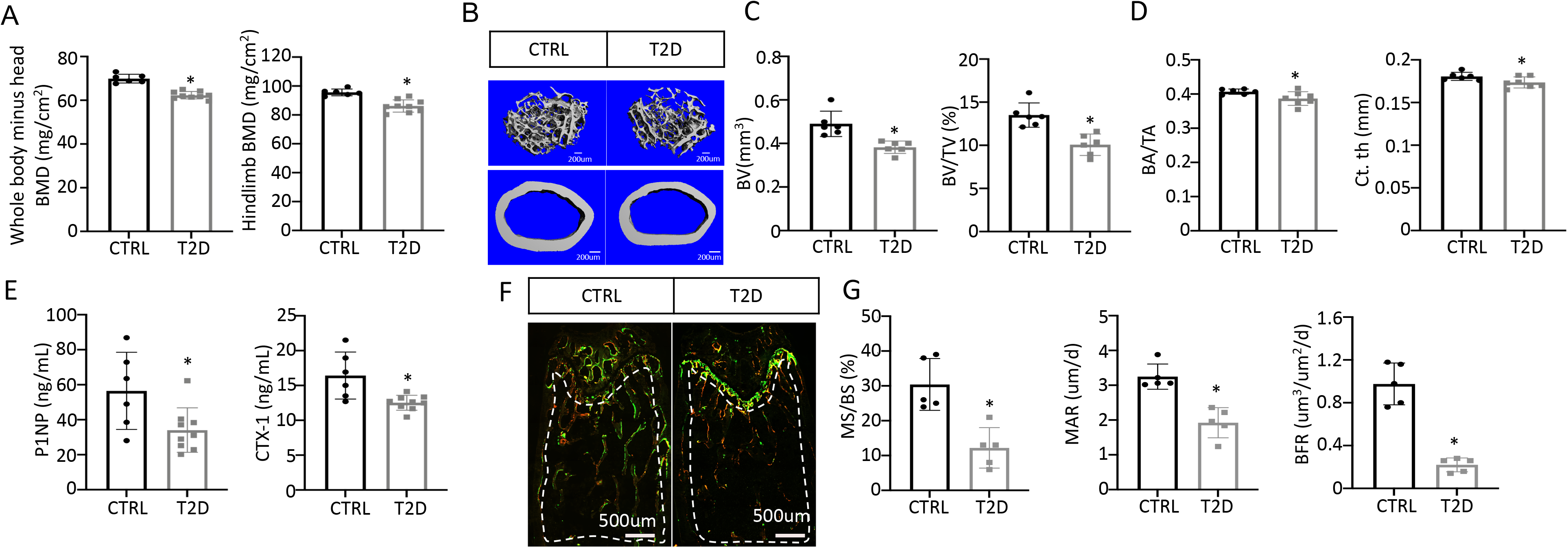
T2D causes low turnover osteopenia. (A) BMD for whole body (minus head) and one leg (femur and tibia). CTRL, n=6; T2D, n=9. (B) Representative µCT images. (C) Quantitative analysis of trabecular bone of distal femur. n=6. (D) Quantitative analysis of cortical bone in femur. CTRL, n=6; T2D, n=7. (E) Serum bone formation (P1NP) and resorption (CTX-1) markers. CTRL, n=6; T2D, n=9. (F, G) Representative images (F) and quantification (G) for double labeling of trabecular bone in distal femur. n=5. Data are mean ± SD. Unpaired Student’s t tests. *: P < 0.05.

We next investigated the cellular basis for diabetic osteopenia. Serum P1NP (procollagen type I N-terminal propeptide) and CTX-1 (collagen type I C-telopeptide) levels were both lower in T2D than control (Fig. 2E), indicating that T2D suppressed the overall bone turnover whereas reduced bone formation was responsible for the net bone loss. To further quantify osteoblast activity, we performed dynamic histomorphometry with dual dyes (calcein followed by alizarin red) that labeled active bone-forming surfaces (Fig. 2F-G, S1J-M). Quantification within the trabecular bone region of the distal femur detected a marked decrease in both mineralizing surface relative to bone surface (MS/BS) and the mineralization apposition rate (MAR), resulting in an 80% suppression of the bone formation rate (BFR/BS) in the T2D mice (Fig. 2F-G). Similarly, MS/BS and BFR/BS were both reduced at the endosteum of the bone shaft whereas MS/BS was also decreased at the periosteum (Fig. S1J-M). These results indicate that T2D impairs osteoblast activity in both trabecular and cortical bone.

### T2D impairs glucose metabolism in bone in vivo

To explore the metabolic basis for impaired osteoblast activity in T2D, we conducted stable isotope tracing in bone with uniformly labeled ^13^C-glucose (^13^C_6_-Glc). Catabolism of ^13^C_6_-Glc was expected to produce fully labeled pyruvate Pyr(m+3) through glycolysis, which could then convert to fully labeled lactate Lac(m+3) in the cytosol or enter mitochondria to produce m+2 labeled TCA cycle metabolites in the first cycle (Fig. 3A). For the experiment, ^13^C_6_-Glc was injected into conscious mice (22 wks of age) via tail vein at 60 mins before the mice were harvested for detection of metabolites in bone or plasma by mass spectrometry. The glycolytic and TCA intermediate pool sizes in both bone (Fig.3B) and plasma (Fig. S2A, B) were similar between T2D and CTRL, while glucose was increased in T2D as expected. The fractional enrichment of ^13^C_6_-Glc in bone reached approximately 20% with no difference between T2D and CTRL mice (Fig. 3C). However, among the bone metabolites detected, the labeling enrichments of Glu(m+2) and Gln(m+2) were significantly reduced in T2D (Fig. 3C). Moreover, when normalized to bone Glc(m+6), the relative enrichments of bone Pyr(m+3), Glu(m+2) and Gln(m+2) were decreased in T2D (Fig. 3D). Further examination of the TCA cycle metabolites in bone revealed substantial enrichment of the m+1 isotopomers, among which Suc(m+1), Asp(m+1), Glu(m+1) and Gln(m+1) were diminished in T2D, with or without normalization to Glc(m+6) in bone (Fig. S3A, B). Therefore, to capture the overall contribution of ^13^C_6_-Glc we calculated the total carbon enrichment ((m0*0+m1*1+m2*2+….mn*n)/n) for each of the TCA cycle metabolites in bone. The result showed that ^13^C enrichments in succinate, glutamate and glutamine, with or without normalization to bone Glc(m+6), were significantly reduced in T2D compared to the CTRL (Fig. 3E). The bone-intrinsic metabolic defect was further supported by Western blot analyses showing a marked decrease in Glut1 and Hk2 in the bone extracts of T2D mice (Fig. 3F, G). Overall, T2D suppresses glucose metabolism in bone through both glycolysis and the TCA cycle.

**Figure 3.**
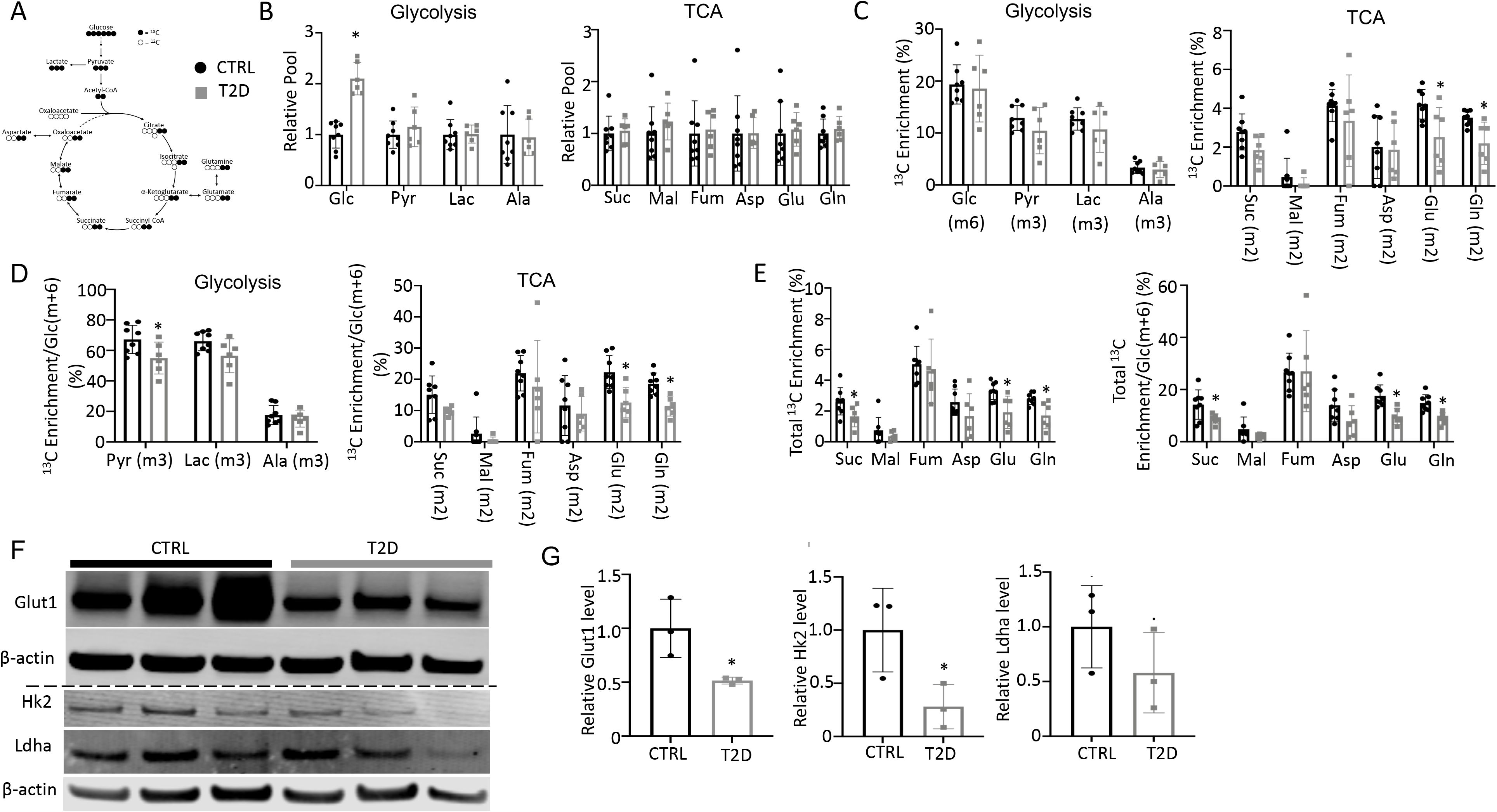
T2D suppresses glucose metabolism in bone. (A) A diagram for carbon tracing with ^13^C_6_-Glc. (B) Relative pool size of glycolytic and TCA metabolites. CTRL, n=8; T2D, n=6. (C) Enrichment of specific isotopologues of metabolites relative to own pool. CTRL, n=8; T2D, n=6. (D) Relative enrichment of specific isotopologues normalized to Glc(m+6). CTRL, n=8; T2D, n=6. (E) Carbon enrichment of metabolites relative to own pool (left) and normalized to Glc(m+6) (right). CTRL, n=8; T2D, n=6. (F, G) Western blot images (F) and quantifications (G). Dashed line denotes two separate gels in F. n=3. Data presented as mean ± SD. *: P < 0.05, Two-tailed unpaired t test.

### T2D suppresses energy metabolism and osteoblast differentiation in bone marrow mesenchymal cells

As the tracing studies above analyzed the bone shaft which contained mostly mature osteoblasts and osteocytes, we next sought to determine whether T2D similarly affected the less mature cells in the osteoblast lineage. To this end, we performed single-cell RNA-sequencing (scRNA-seq) with bone marrow mesenchymal cells purified with FACS from the bone marrow and the digested bone chips of T2D vs CTRL mice (22-week-old). Integrated analyses of the datasets uncovered 20 clusters common to both T2D and control (Fig. 4A). Molecular annotation based on known marker genes identified a majority of the cells as Cxcl12-abundant reticular (CAR) cells including both Adipo-CAR (clusters 1, 2, 5) and Osteo-CAR (cluster 0)^29^ (Fig. 4A, B). Osteoblasts (clusters 3, 15, 18) marked by Bglap and Dmp1 were also readily identified whereas cluster 7 expressed Pdgfra and Ly6a (encoding Sca1) and therefore were likely the PaS mesenchymal progenitors as previously reported^30^. The remaining populations included chondrocytes (cluster 10), pericytes (cluster 16)^31^, endothelial cells (cluster 11), erythrocytes (clusters 4, 6, 8, 19), and several hematopoietic cell types expressing Ptprc (encoding CD45) (clusters 9, 12, 13, 17) together with a minuscule population exhibiting both Osteo-CAR and mature osteoblast markers and therefore an unclear identity (cluster 20). Among CAR cells, osteoblasts, and PaS cells, the relative abundance of CAR cells was slightly increased from 77.6% to 79.2% at the expense of both osteoblasts and PaS cells. Overall, T2D does not overtly alter the composition of bone marrow mesenchymal cells. However, gene set enrichment analysis (GSEA) within individual clusters detected notable differences in metabolic pathways between T2D and CTRL. In particular, T2D suppressed the OXPHOS genes in Adipo-CAR clusters 1 and 5 as well as osteoblast cluster 15, but increased them in Osteo-CAR cluster 0 and osteoblast cluster 3 (Fig. 4C, D) (Supplementary Table S2). In addition, T2D increased glycolysis genes in cluster 3 but suppressed them in cluster 5 (Supplementary Table S2). Thus, T2D appears to disrupt normal energy metabolism among osteoblasts and other mesenchymal cells in the bone marrow.

**Figure 4.**
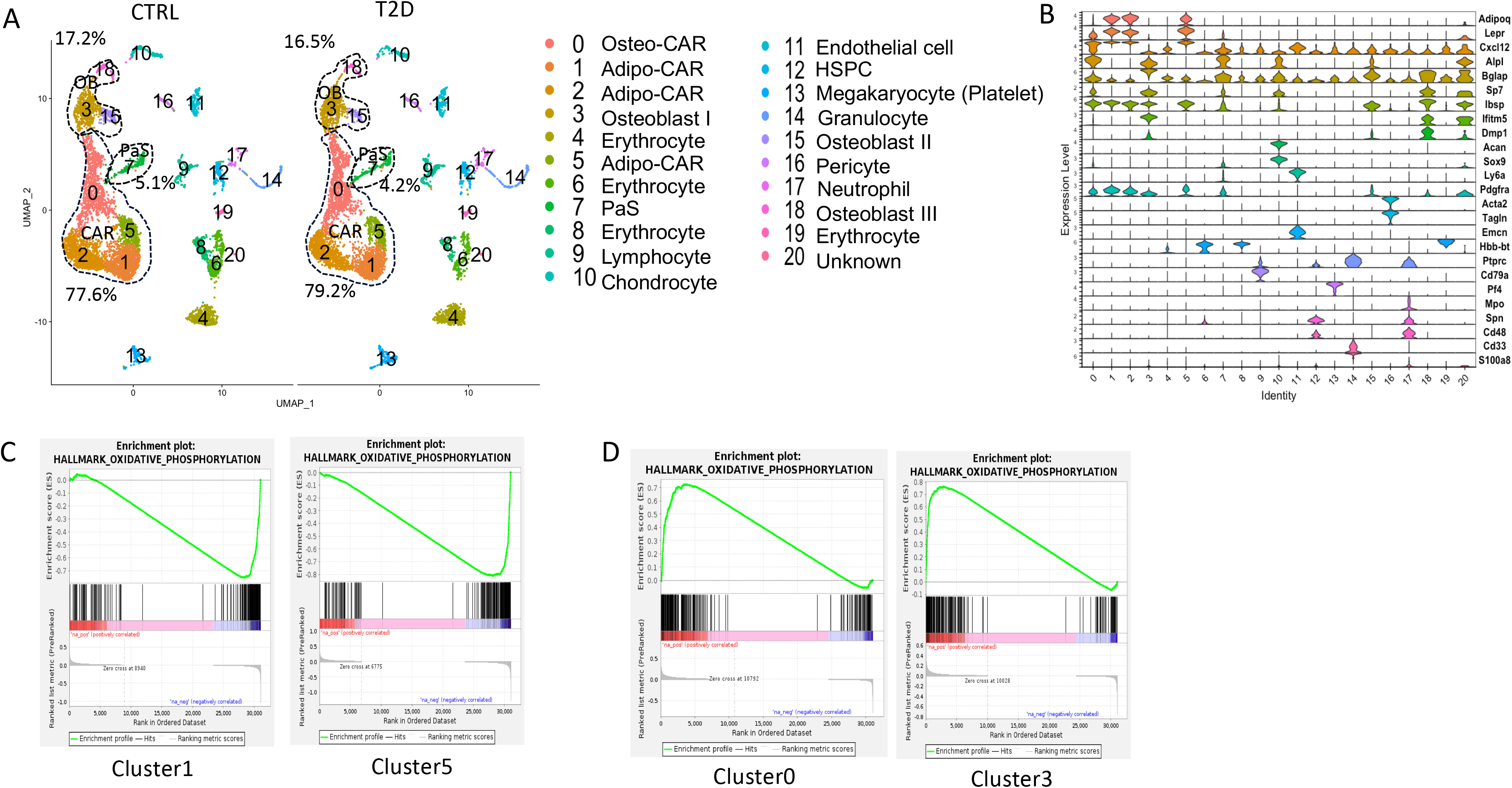
scRNA-seq detects metabolic dysregulation in bone marrow mesenchymal cells of T2D mice. (A) UMAP clusters with annotations to right. Percentages denote relative abundance of osteoblast (OB), CAR and PaS cells among mesenchymal cells. (B) Violin plots of example feature genes for each cluster. (D) GSEA of T2D over CTRL for OXPHOS pathway in clusters as indicated.

To assess the actual changes in energy metabolism, we purified bone marrow stromal cells (BMSC) from the central marrow for metabolic assays in vitro. Immuno-depletion with MACS beads effectively eliminated CD45^+^ hematopoietic cells from the mesenchymal cells (Fig. S4A, B). In the T2D BMSC, both glucose consumption and lactate production were markedly reduced (Fig. 5A). Seahorse assays detected a significant reduction in both oxygen consumption rate (OCR) and extracellular acidification rate (ECAR) in T2D BMSC (Fig. 5B, C). The ATP production rate calculated from either glycolysis (glycoATP) or OXPHOS (mitoATP) was reduced in the T2D cells (Fig. 5D). Thus, T2D suppresses both glycolysis and mitochondrial respiration in BMSC.

**Figure 5.**
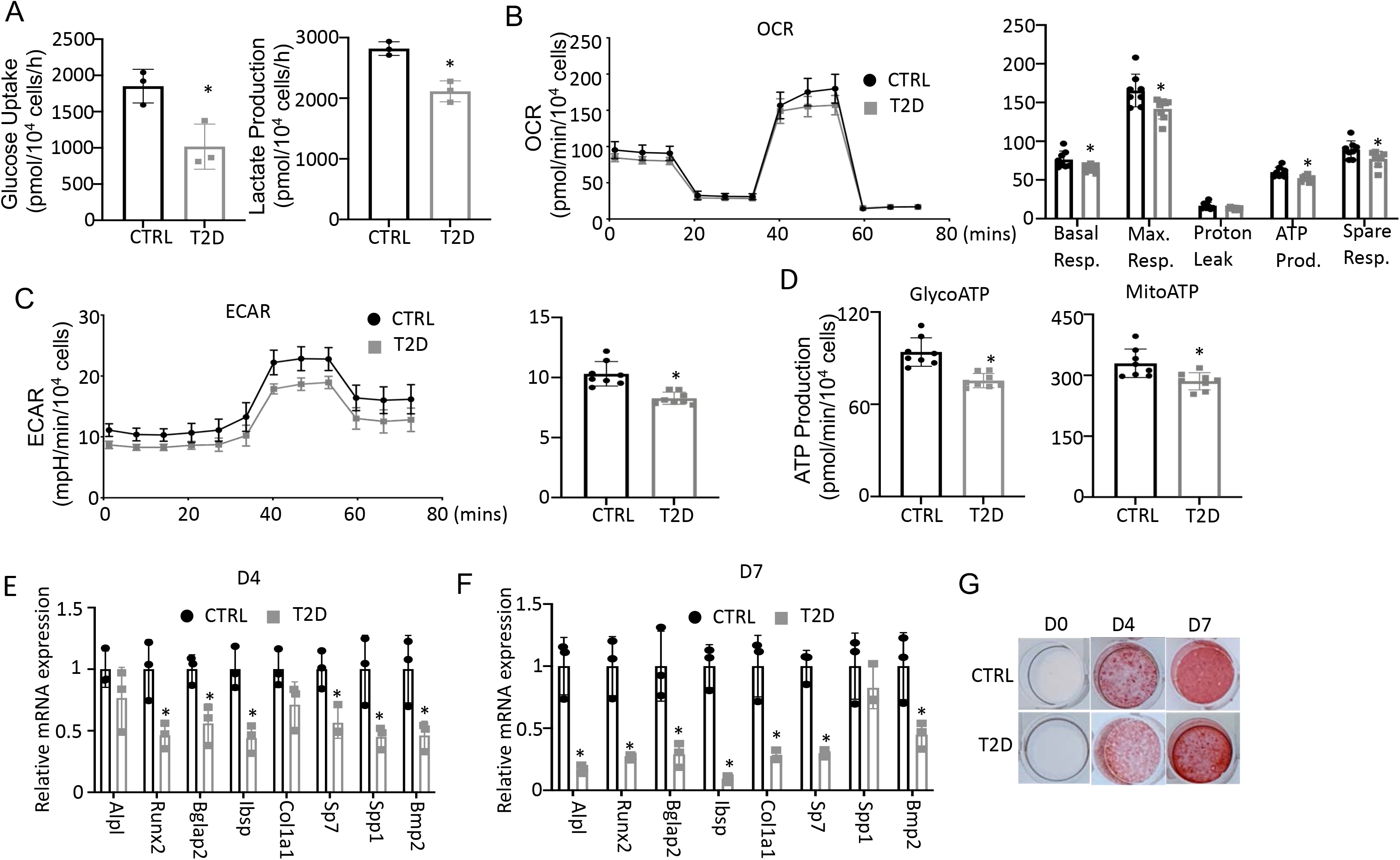
T2D impairs glucose metabolism and osteogenic differentiation in BMSC. (A) Glucose uptake and lactate production rates. n=3. (B, C) Seahorse measurements of OCR (B) and ECAR (C). n=8. (D) Projected ATP production glycolysis (glycoATP) and mitochondria (mitoATP) based on Seahorse. n=8. (E, F) qPCR for osteoblast markers at day 4 (E) and day 7 (F) of differentiation. n=3. (G) Alizarin red staining. Data presented as mean ± SD. *P < 0.05, Unpaired student’s t test.

We next evaluated the potential effect of bioenergetic defects on osteoblast differentiation in T2D. We first confirmed that MACS-purified BMSC underwent robust osteoblast differentiation in vitro. BMSC from normal mice exhibited no Alizarin red staining before differentiation (day 0) but formed intensely stained nodules on day 4 and day 7 (Fig. S5A). qPCR showed that the osteoblast makers Ibsp, Bglap2, Alpl, Spp1, Bmp2, Col1a1, Sp7, and Runx2 were greatly induced after four days of differentiation, with some further upregulated on day 7 (Fig. S5B).

Importantly, when cultured in parallel with the normal cells, T2D BMSC expressed markedly lower levels of virtually all osteoblast markers examined on both day 4 and day 7 of differentiation (Fig. 5E, F). The differentiation defect was further supported by reduced Alizarin red staining of the minerals after 4 or 7 days of differentiation (Figure 5G). Overall, the data indicate that T2D causes cell-intrinsic defects in both metabolic and osteogenic properties of BMSC.

### Enhancing glycolysis with metformin ameliorates bone loss in T2D

To test if impaired glucose utilization drives osteopenia in T2D, we sought to boost glycolysis and examine its effect on bone mass. Metformin is a widely prescribed anti-diabetic and has been proposed to boost glycolysis through a variety of mechanisms^32–35^. We therefore administered metformin to the T2D mice through drinking water and examined the effects on metabolic and bone parameters (Fig. 6A). GTT and ITT showed that glucose tolerance and insulin sensitivity were modestly improved by the metformin regimen although fasting glucose levels remained higher than normal (Fig. 6B-E). Metformin did not affect body weight, body composition or circulating insulin levels in the T2D mice (Fig. S6A-C). Importantly, µCT showed that metformin notably increased trabecular bone mass (BV/TV) mainly due to increased trabecular number (Tb. N) and connectivity (Conn. Dens) coupled with decreased trabecular spacing (Tb. Sp), with no changes in trabecular thickness or cortical bone parameters (Fig. 6F, Fig. S6D, E). Double labeling revealed that metformin increased mineralizing surfaces (MS/BS) without altering the mineral apposition rate (MAR), resulting in a 50% increase in bone formation rate (BFR) in the treated T2D mice (Fig. 6G, H, Supplemental Fig. S6F). Thus, metformin mitigates bone loss in T2D by promoting bone formation.

**Figure 6.**
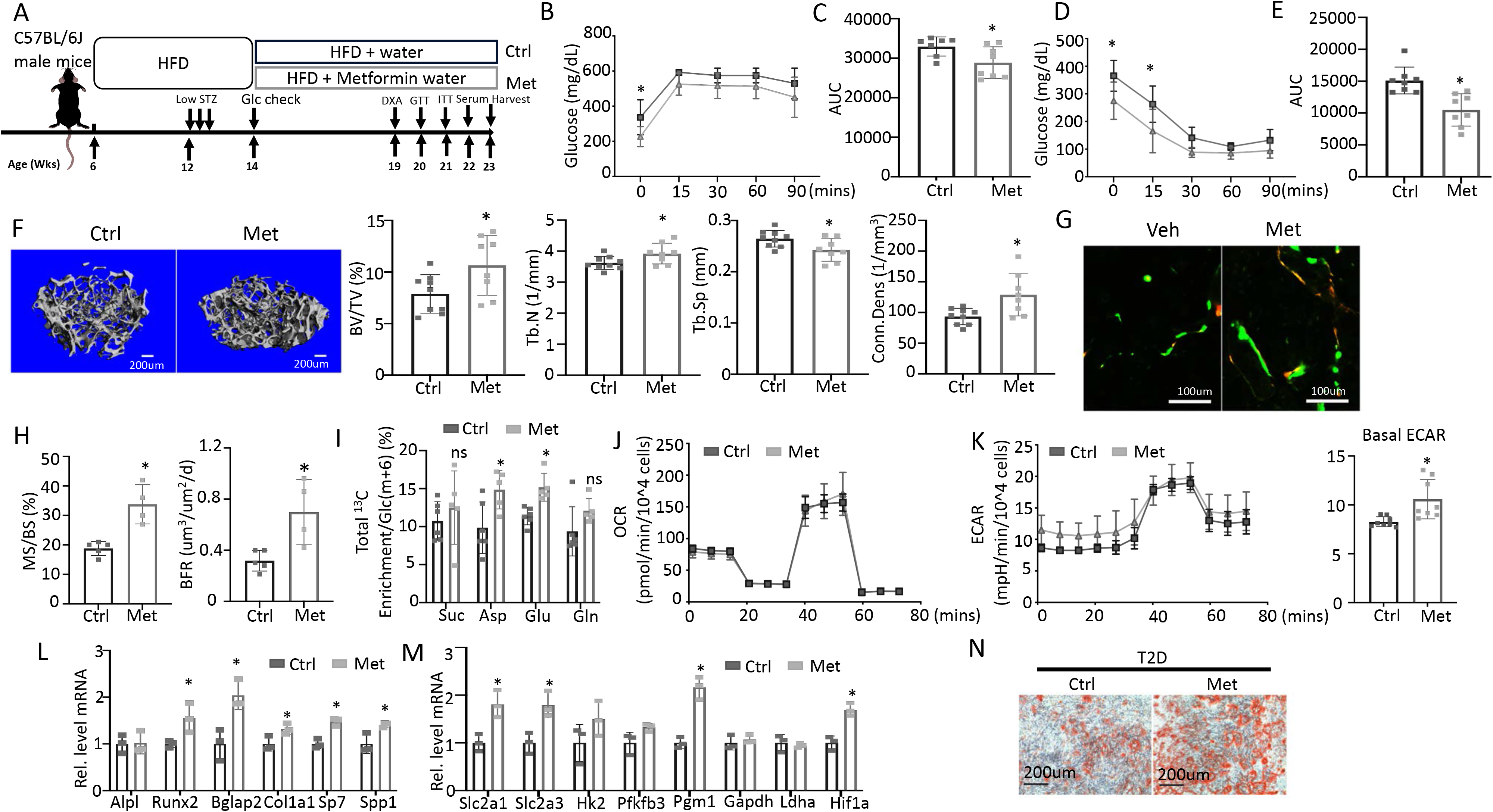
Metformin improves bone mass and glucose metabolism impaired by T2D. (A) A schematic for metformin regimen. T2D animals were randomly divided into Control (Ctrl) and metformin (Met) groups. (B, C) GTT curve (B) and area under curve (C). Ctrl, n=7; Met, n=8. (D, E) ITT curve (D) and area under curve (E). n=8. (F) Representative uCT images and quantification of trabecular bone in distal femur. Ctrl, n=9; Met, n=8. (G, H) Representative images (G) and quantification (H) of double labeling in trabecular bone of distal femur. Ctrl, n=5; Met, n=4. (I) Carbon enrichment of TCA metabolites normalized to Glc(m+6). Ctrl, n=6; Met, n=5. (J, K) Seahorse measurements of OCR (J) and ECAR (K) with or without 1 mM metformin treatment for 24 hrs. n=8. (L, M) qPCR analyses of osteoblast markers (L) and glycolysis-related genes (M) in BMSC at day 7 of differentiation with or without 1 mM metformin treatment. n=3. (N) Alizarin red staining at day 7 of osteoblast differentiation. Data are presented as mean ± SD. *: P < 0.05, two-way ANOVA followed by Sidak’s multiple comparisons (B, D) or Student’s t test (all others).

To assess the potential effect of metformin on glucose metabolism in bone, we next performed stable isotope tracing with ^13^C_6_-Glc in the T2D mice with or without metformin treatment immediately before harvest at 23 weeks of age (Fig. 6A). Both Pyr(m+3) and Lac(m+3) normalized to Glc(m+6) in bone exhibited a trend of increase in response to metformin, but the differences did not achieve statistical significance, likely due to insufficient power of the sample size (Fig. S6G p=0.05). On the other hand, the total carbon enrichment of aspartate and glutamate relative to Glc(m+6) was significantly increased by metformin, indicating increased glucose entry into the TCA cycle in bone (Fig. 6I). The data, therefore, support that metformin promotes bone glucose metabolism in T2D mice.

We next examined whether metformin has a direct effect on osteoblast-lineage cells. For metabolic effects, BMSC from T2D mice were treated with metformin for 24 hours before Seahorse assays. Metformin had negligible effect on OCR but significantly elevated ECAR (Fig. 6J, K). Moreover, in osteoblast differentiation assays, metformin increased the expression of multiple osteoblast markers and key glycolysis genes in T2D BMSC (Fig. 6L, M). Similarly, metformin notably improved the formation of mineralized nodules by the T2D cells following osteoblast differentiation (Fig. 6N). Together, these findings indicate that metformin directly enhances both glycolysis and osteoblast differentiation in T2D BMSC.

### Genetic activation of glycolysis reduces bone loss in T2D

To corroborate the direct effect on bone, we utilized a transgenic mouse model that increased glycolysis specifically in the osteoblast-lineage cells. The transcription factor Hif1α is known to stimulate the expression of numerous genes in the glycolytic pathway, thus promoting glycolysis. Here, we generated mice with the genotype of Osx-rtTA; tetO-Cre; Hif1dPA (Hif1OE hereafter) to allow for Cre-mediated expression of a degradation-resistant form of Hif1a (Hif1dPA) in the osteoblast lineage upon Dox administration. Their sex-matched littermates missing at least one of the three transgenes were used as controls (Ctrl). Both Hif1OE and Ctrl males were subjected to the established protocol to develop T2D before Dox was supplied in their drinking water until harvest (Fig. 7A). Gene expression studies confirmed that Hif1a and several glycolysis-related genes including Pfkfb3, Ldha and Ldhb, were upregulated in Hif1OE over Ctrl mice (Fig. 7B). µCT 3D reconstruction images revealed a marked increase in both trabecular bone and cortical bone of Hif1OE over Ctrl mice (Fig. 7C). Quantification confirmed that the trabecular bone volume (BV) and bone volume fraction (BV/TV), together with the cortical bone area (BA), bone area fraction (BA/TA) and thickness (Ct. Th) were all increased in Hif1OE (Fig. 7D, Fig. S7A, B). Double labeling experiments detected notable increases in MS/BS and BFR/BS but not MAR in trabecular bone (Fig. 7E, Fig. S7C). Overall, Hif1a overexpression in osteoblast-lineage cells increases bone mass in T2D mice.

**Figure 7.**
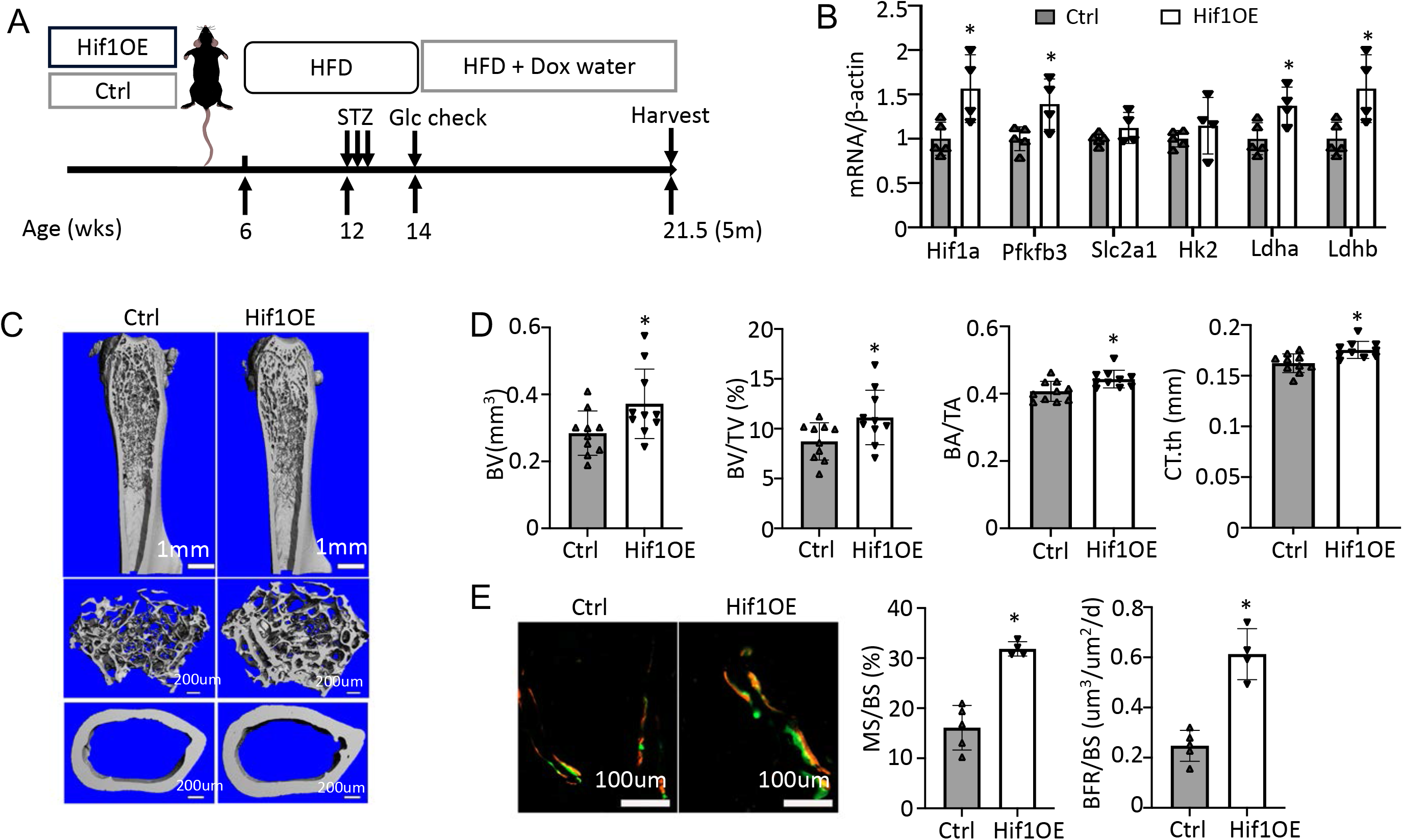
Targeted overexpression of Hif1a in osteoblasts rescues low bone mass in T2D mice. (A) A schematic for Hif1a overexpression (Hif1OE). (B) qPCR analyses of glycolysis related genes in bone. n=4. (C, D) Representative uCT images (C) and quantification of distal trabecular bone and cortical bone (D) in femur. n=10. (E) Representative images and quantification of double labeling in trabecular bone of proximal tibia. Ctrl, n=5, Hif1OE, n=4. Data presented as mean ± SD. *: P < 0.05 Unpaired Student’s t test.

We further investigated the rate-determining steps in the glycolytic pathway, which could be therapeutically targeted for promoting bone formation in T2D mice. Overexpression of Glut1, a major glucose transporter in osteoblasts, has been shown to promote bone formation in normal mice^36^. We, therefore, generated mice with the genotype of Osx-rtTA; TRE-Glut1 (Glut1OE) to induce Glut1 overexpression in osteoblast-lineage cells with Dox. Both Glut1OE males and their sex-matched littermates (Ctrl) missing one of the two transgenes were rendered diabetic and then exposed to Dox according to the same regimen as described for the Hif1OE mice. Western blot analyses demonstrated successful overexpression of Glut1 in the bone shaft of long bones in Glut1OE (Fig. S8A). However, µCT analysis detected no significant change in either trabecular or cortical bone parameters between Glut1OE and Ctrl mice (Fig. S8B, C). Likewise, serum P1NP and CTX-1 levels were similar between the genotypes (Fig. S8D). Thus, increased Glut1 expression does not improve osteopenia in T2D mice.

We next tested the potential effect of phosphofructokinase-2/fructose-2,6-bisphosphatase (Pfkfb3). Pfkfb3 synthesizes fructose-2,6-bisphosphate (F2,6P2), a potent allosteric activator of 6-phosphofructo-1-kinase (Pfk1) which controls a rate-limiting step of glycolysis^37^. To this end, we generated Osx-rtTA; TRE-Pfkfb3 mice (Pfkfb3OE) along with their control littermates (Ctrl) missing at least one of the two transgenes. Cohorts of Pfkfb3OE and Ctrl mice were each divided into a normal or T2D group. In the T2D group, the mice were rendered diabetic and then treated with Dox for the remainder of the experiment, whereas those in the normal group were maintained on regular chow before being treated with Dox in the same way (Fig. 8A). Upon harvest of the mice, µCT analyses detected no difference in any of the trabecular bone parameters between Pfkfb3OE and Ctrl mice in the normal group (Fig. 8B, C, Fig. S9A). However, within the T2D group, Pfkfb3OE significantly increased trabecular bone volume (BV), trabecular bone fraction (BV/TV), coupled with increased connectivity density (Conn. Dens), increased trabecular number (Tb. N) and reduced trabecular spacing (Tb. Sp) (Fig. 8B, C, Fig. S9A). Similarly, Pfkfb3 overexpression did not change the cortical bone parameters in the normal group, but it increased cortical bone fraction (BA/TA) in the T2D group, thanks to the correction of total cross-sectional area (TA) (Fig. 8D, Fig. S9B). Dynamic histomorphometry revealed that Pfkfb3 overexpression in T2D mice increased bone formation rate (BFR) due to a marked increase in bone mineralizing surface (MS/BS) without affecting the mineral apposition rate (MAR) (Fig. 8E, Fig. S9C). Serum biochemical assays showed that Pfkfb3 overexpression elevated P1NP levels without changing CTX-I in the T2D mice (Fig. 8F). RT-qPCR detected a modest increase of Pfkfb3, Hif1a, Glut1, and Ldhb mRNA in the bone shaft of Pfkfb3OE diabetic mice versus Ctrl T2D mice (Fig. 8G). Overall, the results demonstrate that bone-specific activation of glycolysis is an effective anabolic strategy to counter the bone loss in T2D mice.

**Figure 8.**
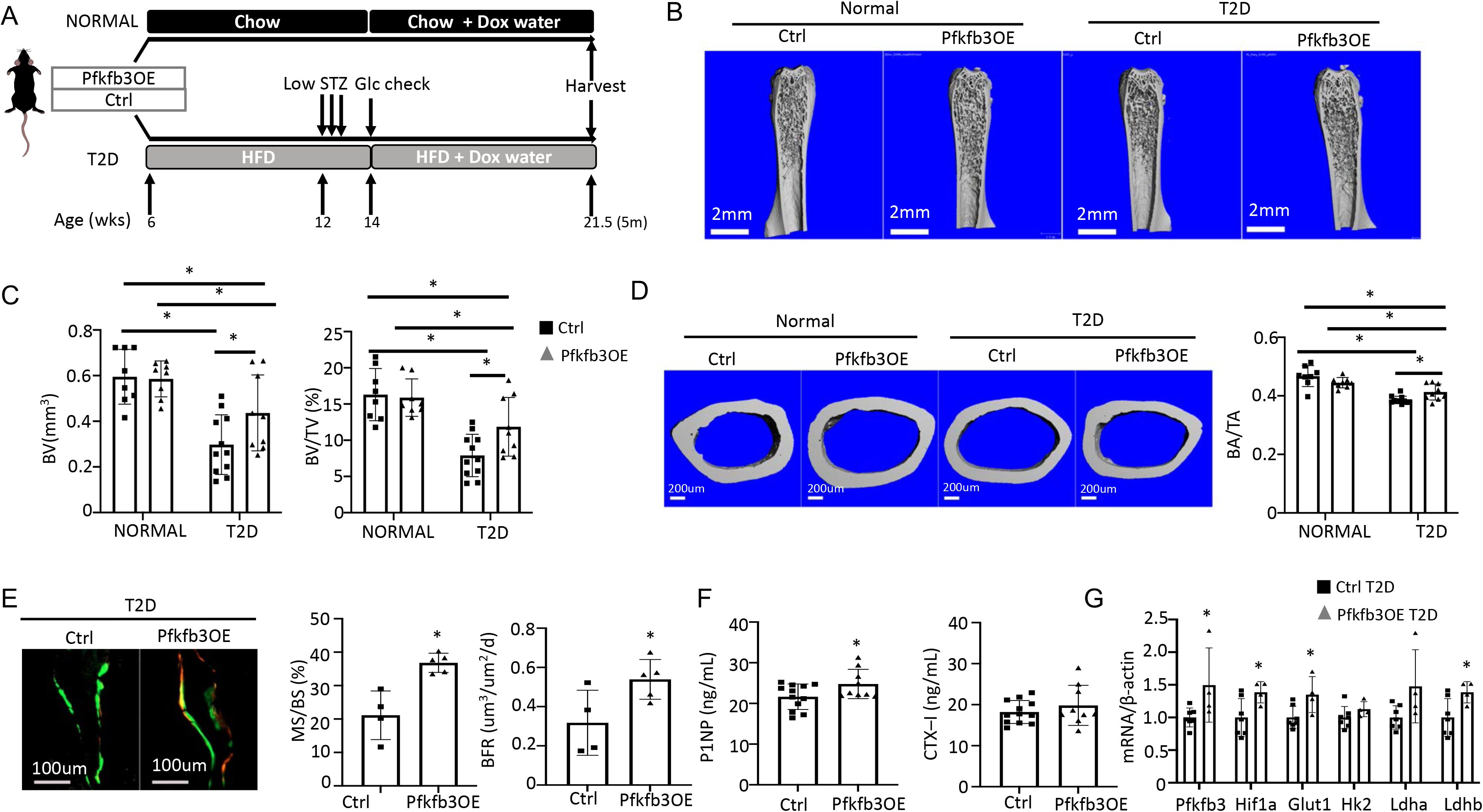
Osteoblast-directed Pfkfb3 overexpression corrects bone loss in T2D. (A) A schematic for experimental design. Representative uCT images (B) and quantification (C) of trabecular bone in distal femur. (D) Representative uCT images and quantification of cortical bone in femur. Ctrl NORMAL, n = 8; Pfkfb3OE NORMAL, n = 8; Ctrl T2D, n = 11; Pfkfb3OE T2D, n = 9. (E) Representative images and quantification of double labeling in trabecular bone of proximal tibia. Ctrl T2D, n = 4; Pfkfb3OE T2D, n = 5. (F) Serum markers for bone formation (P1NP) and resorption (CTX-1). Ctrl T2D, n = 12; Pfkfb3OE T2D, n = 9. (G) qPCR analyses of glycolysis related genes in bone. Ctrl T2D, n = 7; Pfkfb3OE T2D n = 4. Data presented as mean ± SD. *: P < 0.05, Two-way ANOVA followed by Fisher’s LSD test. (C, D) or Student’s t test (all others).

## Discussion

We have investigated the mechanism underlying the bone anabolic defect in a T2D mouse model. Evidence from multiple approaches including stable isotope tracing in vivo, scRNA-seq of bone marrow mesenchymal cells, as well as in vitro flux assays supports that T2D causes metabolic dysregulation intrinsic to osteoblast lineage cells. Importantly, activation of glycolysis either pharmacologically or by genetic means effectively enhances bone formation and preserves bone mass in diabetic mice. The results therefore support osteoblast-intrinsic impairment of glucose metabolism as a pathogenic mechanism for diabetic osteopenia in T2D.

Although adult-onset T2D differs from youth-onset T2D in that the net bone mass is either normal or even increased in adults, they both suppress bone formation. Multiple mechanisms including hyperglycemia and decreased signaling by Wnt, adipokine, insulin, or Igf1 have been proposed to impair osteoblast differentiation and function in diabetes, but their functional relevance in vivo remains elusive^38^. Nonetheless, as Wnt and Igf1 signaling have been shown to promote glycolysis directly in osteoblast lineage cells, it is reasonable to speculate that downregulation of the aforementioned signals may be a proximate cause for impaired glycolysis in diabetic bone^6, 7^. Our mouse model recapitulates the low bone turnover as seen in human patients, but we have chosen to focus on the bone anabolic defect as it drives the osteopenia phenotype. The model can be further studied in the future to determine the mechanism for impaired bone resorption in T2D and its potential contribution to impaired bone formation through a coupling mechanism. Nonetheless, it is worth noting that osteoblast-directed activation of glycolysis increased bone formation activity without any “coupled” activation of bone resorption, indicating that osteoblasts can be selectively targeted to increase bone mass in T2D.

Our data support the functional contribution of osteoblast glucose metabolism to the bone anabolic defect in T2D. Metformin notably increased bone formation in the T2D mice, concurrent with stimulation of glucose metabolism in bone. Similarly, in vitro, metformin enhanced glycolysis together with osteoblast differentiation in BMSC derived from T2D mice, indicating a direct effect on osteoblast lineage cells. Targeted overexpression of Hif1a, known to activate multiple target genes in the glycolysis pathway, markedly increased bone formation and restored bone mass in T2D mice. Finally, overexpression of a single glycolysis-promoting gene Pfkfb3 was sufficient to overcome the bone anabolic defect caused by T2D. In contrast, overexpression of Glut1 failed to increase bone formation, indicating that glucose uptake was not responsible for the reduced glycolysis flux or impaired bone anabolic activity in T2D. The result further supports the notion that downregulation of Glut1 in diabetic bones likely represents a protective mechanism against cellular damages, as intracellular accumulation of glucose is known to be cytotoxic through multiple mechanisms^39^.

In conclusion, the study identifies impaired glucose metabolism in osteoblasts as a key mediator for the bone anabolic defect in T2D. Furthermore, osteoblast glycolysis offers a promising target for developing future bone therapies.

## Acknowledgements

The work is partially supported by NIH grant R01 DK125498 (FL).

## Data availability

ScRNA-seq data have been deposited into GEO repository with accession number GSE221936. All data related to the figures have been uploaded to Dryad (doi:10.5061/dryad.s7h44j1bc).

## Supplemental figure legends

**Figure S1.**
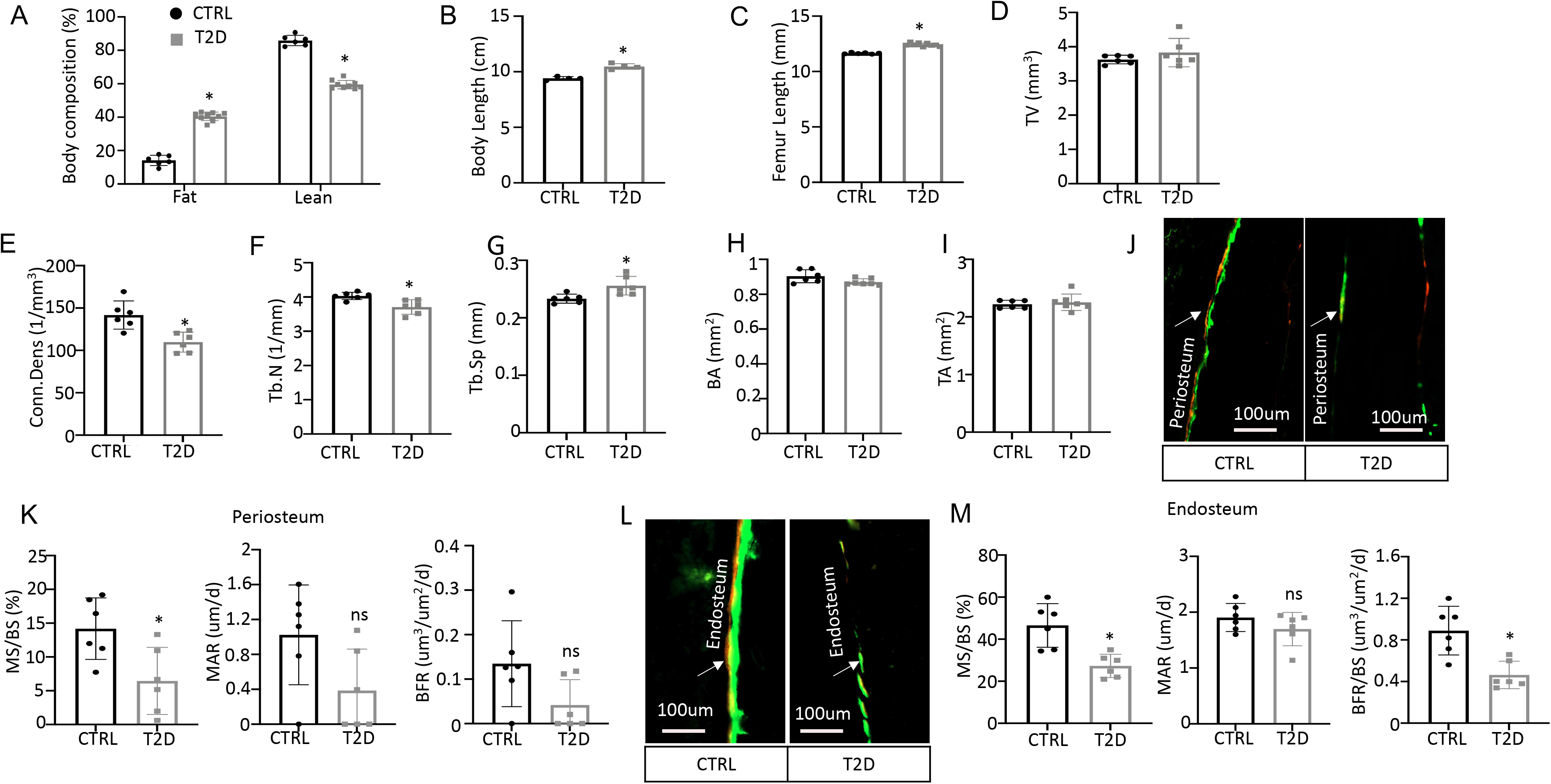
Diabetic osteopenia in T2D mouse model. (A) Body composition. CTRL, n=6; T2D, n=9. (B) Body length (cm) from nose tip to anus. n=4. (C) Femur length measured by DXA. CTRL, n=6; T2D, n=9. (D-G) Trabecular bone parameters by µCT. n=6. (H) Cortical bone parameters by µCT. CTRL, n=6; T2D, n=7. (J-K) Double labeling analysis of periosteum. n=6. (L-M) Double labeling analysis of endosteum. n=6. Data are represented as mean ± SD. *P < 0.05 as determined by Student’s t test between two groups.

**Figure S2.**
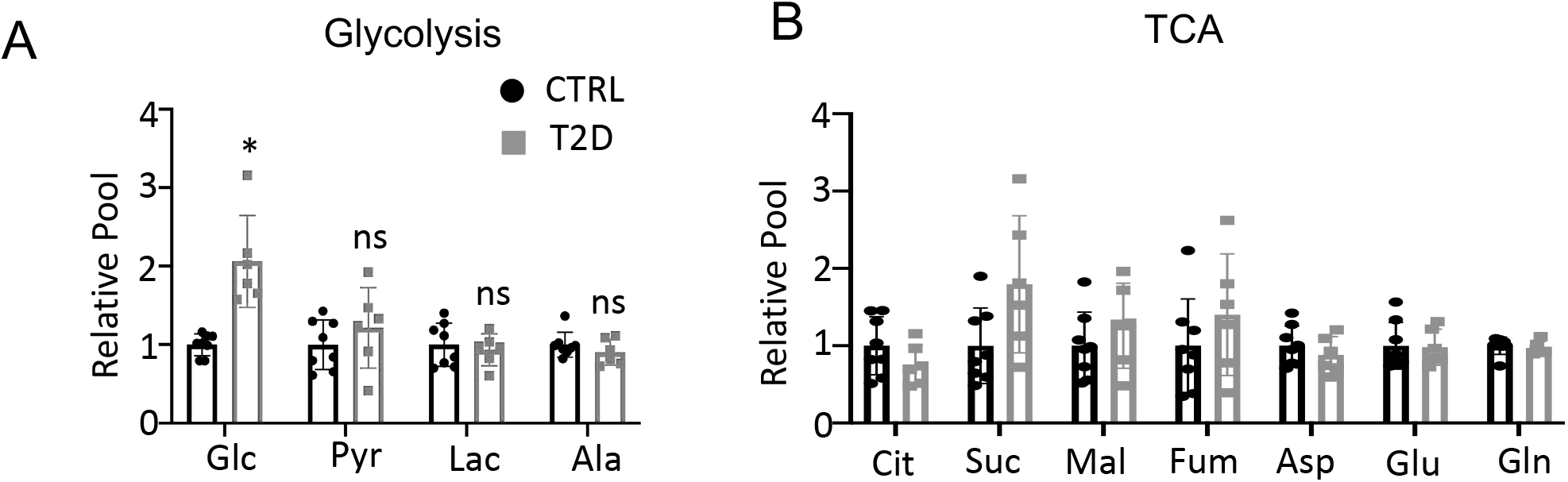
Glucose tracing in plasma. (A, B) Relative levels of glycolysis (A) and TCA (B) metabolites in the plasma. CTRL, n=8; T2D, n=6. Data are represented as mean ± SD. *P < 0.05, Student’s t test.

**Figure S3.**
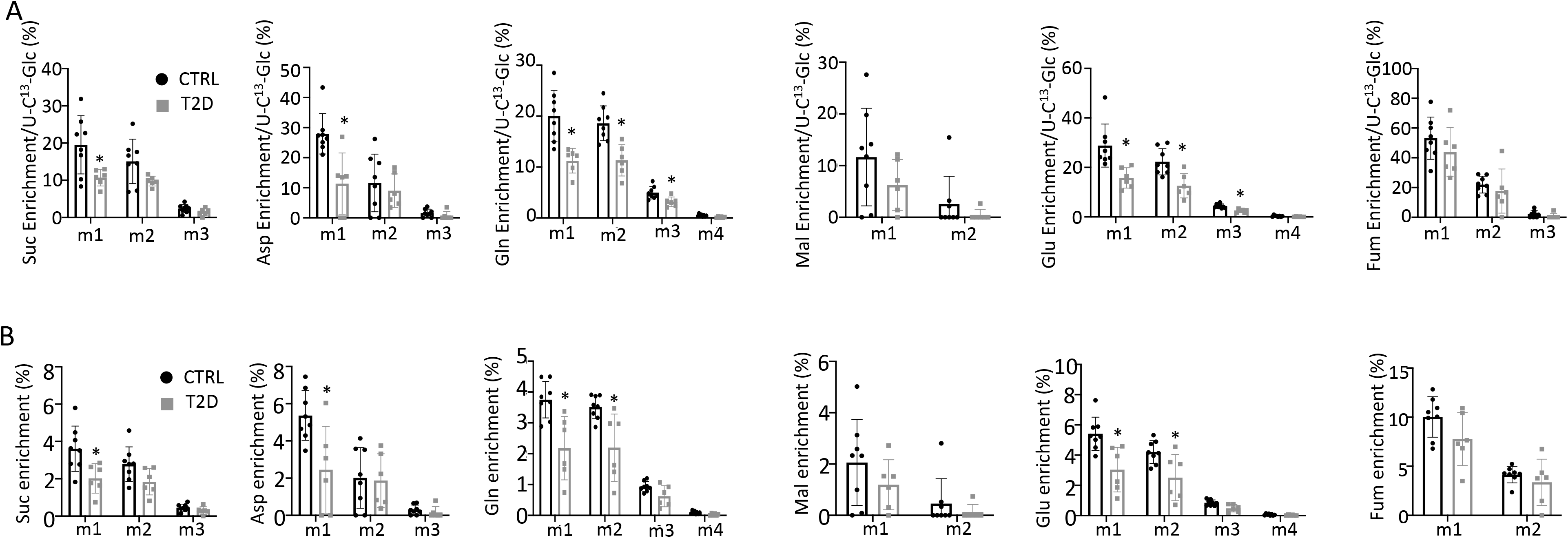
Glucose tracing in bone. (A) Relative enrichment of specific isotopomers normalized to ^13^C_6_-Glc in bone. (B) Enrichment of specific isopotomers in bone. CTRL, n=8; T2D, n=6. Data are represented as mean ± SD. *P < 0.05, two-way ANOVA followed by Sidak’s multiple comparisons.

**Figure S4.**
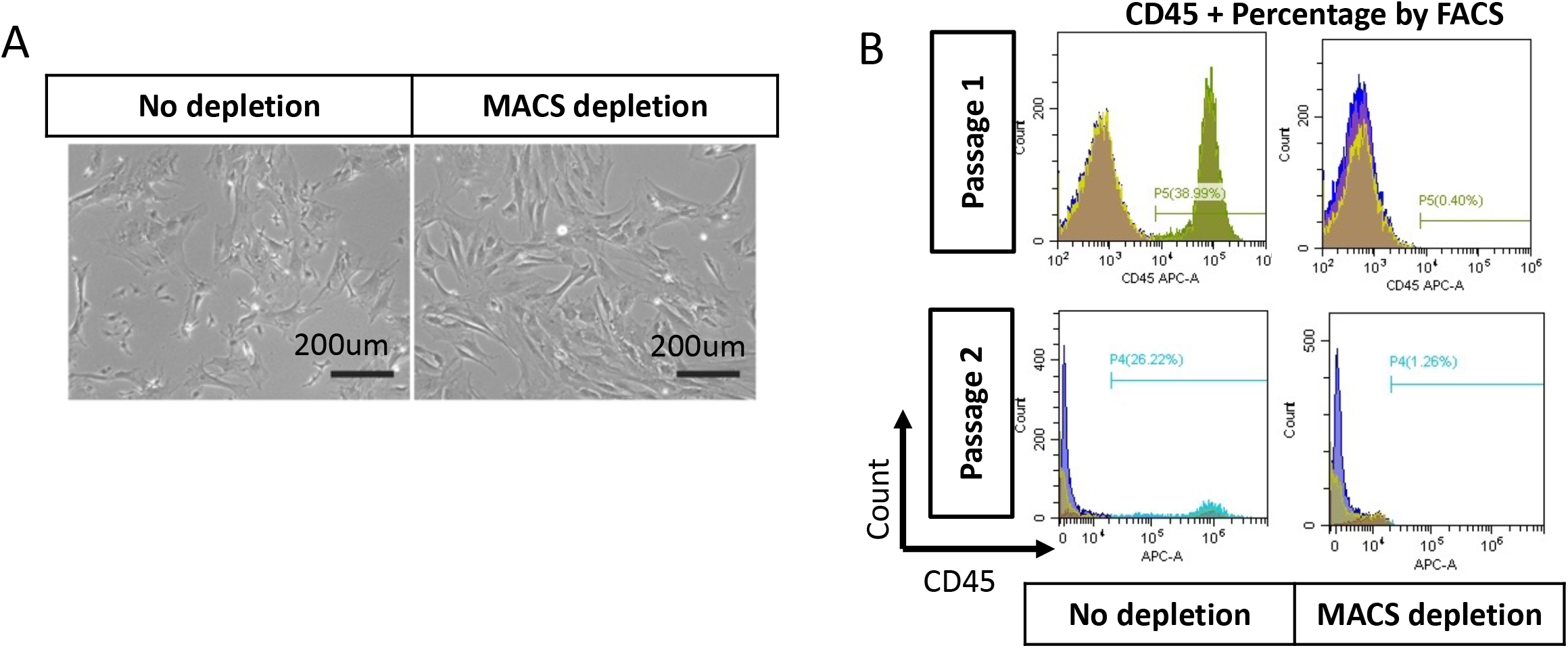
BMSC purification with MACS beads. (A) Bright field of BMSC with or without depletion of CD45+ cells. (B) FACS analysis of CD45^+^ cell percentage for BMSC with or without depletion after one or two passages.

**Figure S5.**
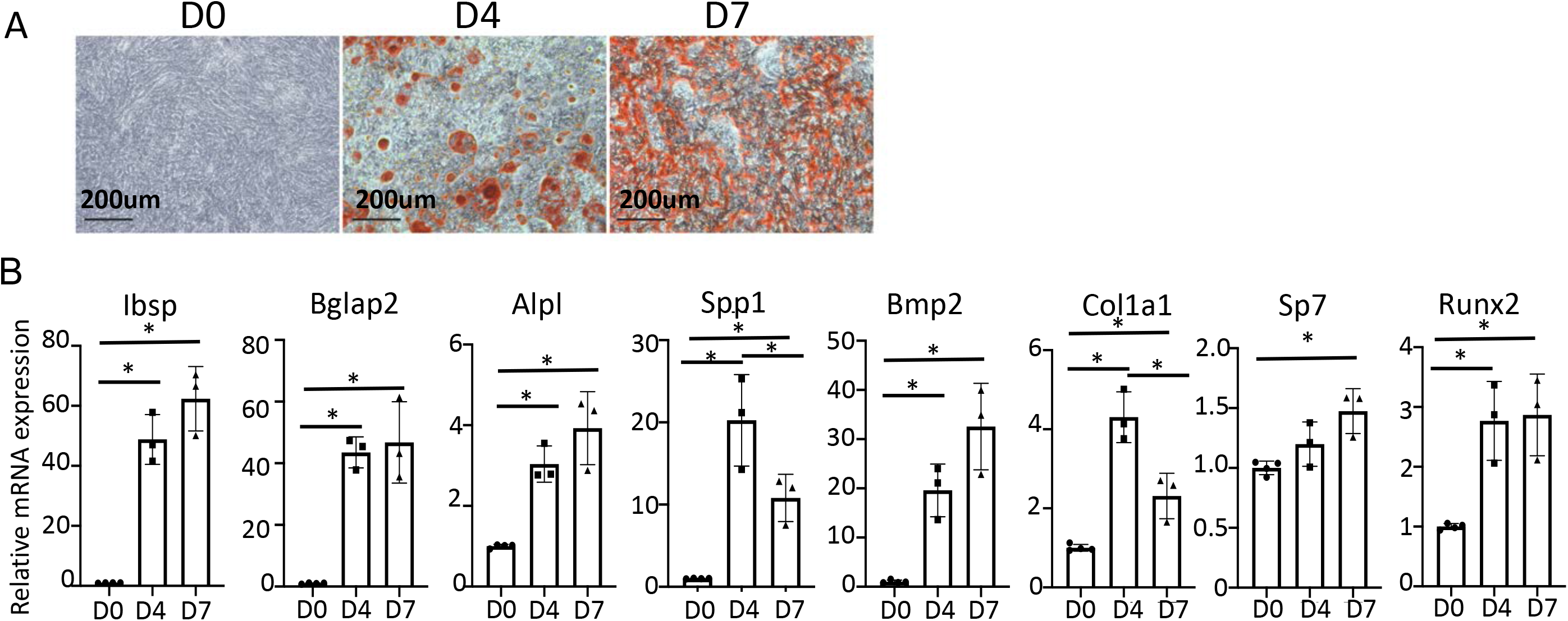
In vitro osteoblast differentiation of MACS-purified BMSC from wild type mice. (A) Alizarin red staining with differentiation at day 0, day 4 and day 7. (B) qPCR of osteoblast marker genes. n=3. Data are represented as mean ± SD. *P < 0.05 by one-way ANOVA followed by Student’s t test.

**Figure S6.**
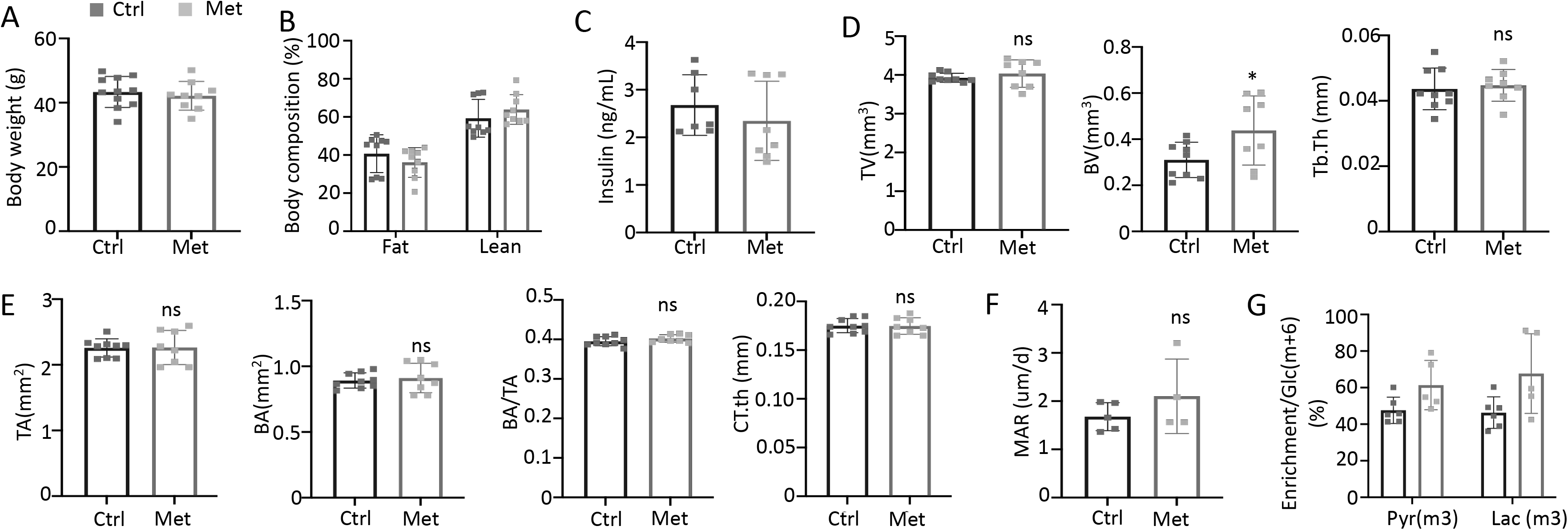
Metformin improves bone mass in T2D mice. (A) Body weight. Ctrl, n=11; Met, n=9. (B) Body composition. n=9. (C) Serum insulin level. Ctrl, n=7; Met, n=8. (D) Trabecular bone parameters of femurs by µCT. Ctrl, n=9; Met, n=8. (E) Cortical bone parameters of femur midshaft. Ctrl, n=9; Met, n=8. (F) Mineral apposition rate by double labeling method. Ctrl, n=5; Met, n=4. (G) Relative enrichment of specific isotopomers normalized to Glc(m+6). Ctrl, n=6; Met, n=5. Data are represented as mean ± SD. *P < 0.05 by Student’s t test.

**Figure S7.**
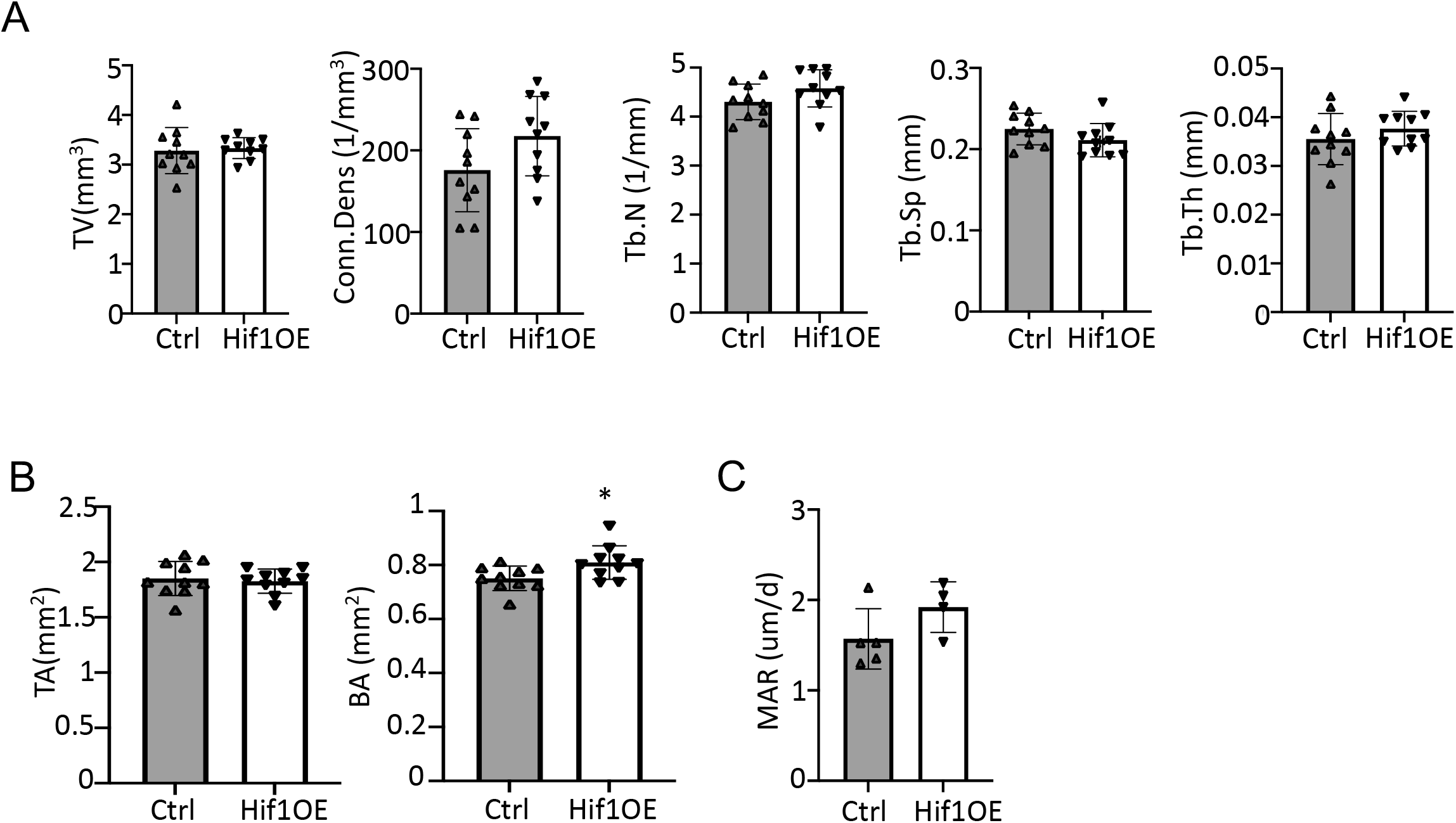
Hif1a overexpression improves bone mass in T2D mice. (A, B) Trabecular (A) and cortical (B) bone parameters by µCT. n=10. (C) Mineral apposition rate (MAR) from double labeling. Ctrl, n=5; Hif1OE; n=4. Data are represented as mean ± SD. *P < 0.05, Student’s t test.

**Figure S8.**
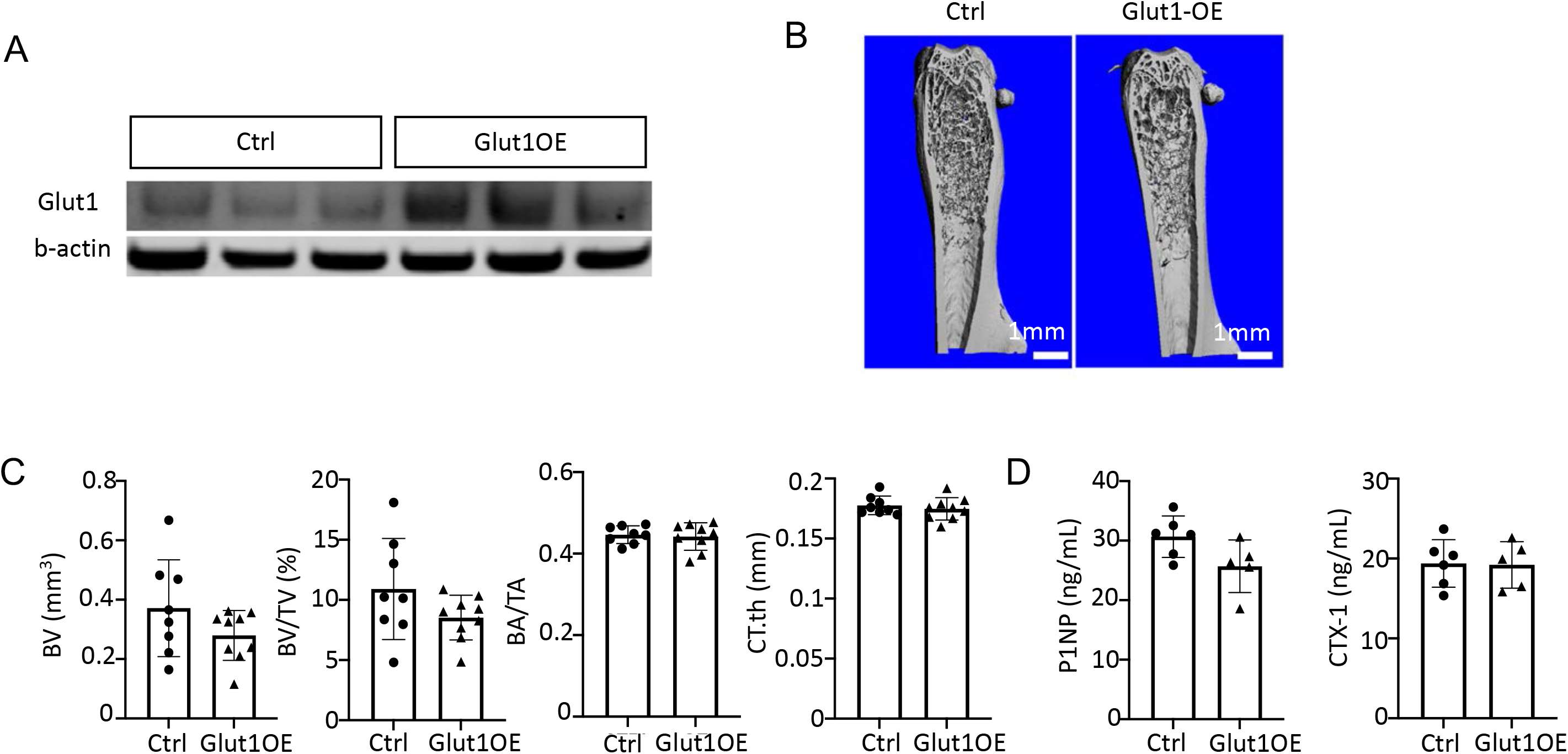
Glut1 overexpression does not improve bone in T2D mice. (A) Glut1 overexpression in bone shaft by Western blot. (B) Representative µCT images. (C) Quantification by µCT. Ctrl, n=8; Glut1OE; n=9. (D) Serum P1NP and CTX-1 levels. Ctrl, n=6; Glut1OE; n=5. Data are represented as mean ± SD. *P < 0.05 by Student’s t test.

**Figure S9.**
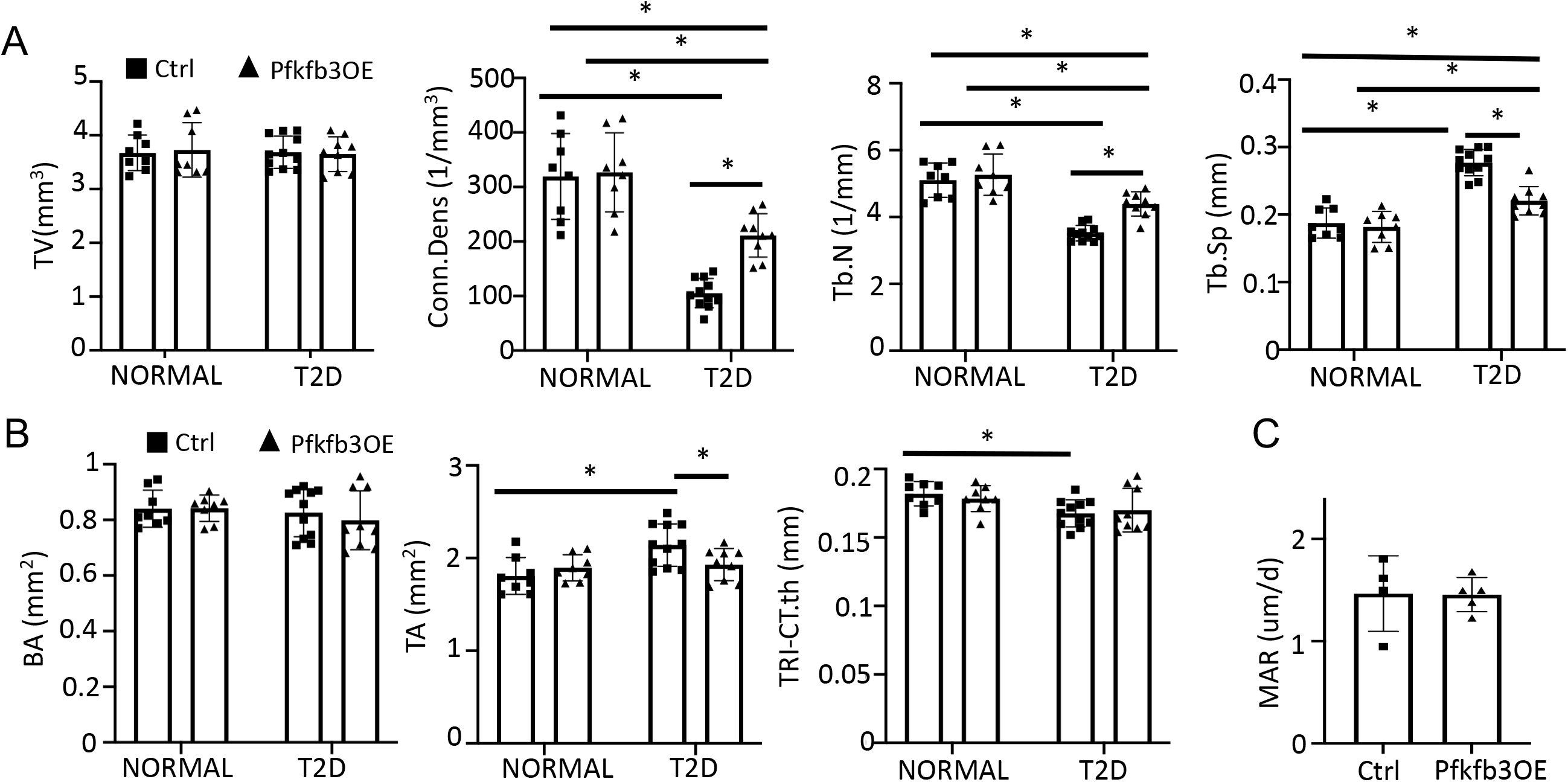
Pfkfb3 overexpression improves bone mass in T2D mice. (A, B) Trabecular (A) and cortical (B) bone parameters of Pfkfb3OE versus Ctrl mice in T2D or NORMAL groups. Ctrl NORMAL, n = 8; Pfkfb3OE NORMAL, n = 8; Ctrl T2D, n = 11; Pfkfb3OE T2D, n = 9. (C) Quantification of mineral apposition rate (MAR) by double labeling in T2D mice with or without Pfkfb3OE. Ctrl T2D, n = 4; Pfkfb3OE T2D, n = 5. Data are represented as mean ± SD. *P < 0.05 Two-way ANOVA followed by Fisher’s LSD test (A, B) or by Student’s t test (C).

## Supplemental table captions

**Table S1 Primer sequences for qPCR.**

**Table S2 GSEA analysis of each cluster (T2D VS CTRL) (FDR q<0.25)**

## References

1. Eriksen, E. F. Cellular mechanisms of bone remodeling. Rev Endocr Metab Disord 11, 219–227, doi:10.1007/s11154-010-9153-1 (2010).

2. Long, F. Building strong bones: molecular regulation of the osteoblast lineage. Nature reviews. Molecular cell biology 13, 27–38, doi:10.1038/nrm3254 (2012).

3. Buttgereit, F. & Brand, M. D. A hierarchy of ATP-consuming processes in mammalian cells. Biochem J 312 (Pt 1), 163–167, doi:10.1042/bj3120163 (1995).

4. Rolfe, D. F. & Brown, G. C. Cellular energy utilization and molecular origin of standard metabolic rate in mammals. Physiol Rev 77, 731–758, doi:10.1152/physrev.1997.77.3.731 (1997).

5. Lee, W. C., Guntur, A. R., Long, F. & Rosen, C. J. Energy Metabolism of the Osteoblast: Implications for Osteoporosis. Endocr Rev, doi:10.1210/er.2017-00064 (2017).

6. Esen, E. et al. WNT-LRP5 signaling induces Warburg effect through mTORC2 activation during osteoblast differentiation. Cell metabolism 17, 745–755, doi:10.1016/j.cmet.2013.03.017 (2013).

7. Esen, E., Lee, S. Y., Wice, B. M. & Long, F. PTH Promotes Bone Anabolism by Stimulating Aerobic Glycolysis Via IGF Signaling. Journal of bone and mineral research : the official journal of the American Society for Bone and Mineral Research, doi:10.1002/jbmr.2556 (2015).

8. Chen, H. et al. Increased glycolysis mediates Wnt7b-induced bone formation. FASEB J 33, 7810–7821, doi:10.1096/fj.201900201RR (2019).

9. Regan, J. N. et al. Up-regulation of glycolytic metabolism is required for HIF1alpha-driven bone formation. Proceedings of the National Academy of Sciences of the United States of America 111, 8673–8678, doi:10.1073/pnas.1324290111 (2014).

10. Lee, S. Y. & Long, F. Notch signaling suppresses glucose metabolism in mesenchymal progenitors to restrict osteoblast differentiation. J Clin Invest 128, 5573–5586, doi:10.1172/JCI96221 (2018).

11. Wei, J. et al. Glucose Uptake and Runx2 Synergize to Orchestrate Osteoblast Differentiation and Bone Formation. Cell 161, 1576–1591, doi:10.1016/j.cell.2015.05.029 (2015).

12. Zoch, M. L., Abou, D. S., Clemens, T. L., Thorek, D. L. & Riddle, R. C. In vivo radiometric analysis of glucose uptake and distribution in mouse bone. Bone Res 4, 16004, doi:10.1038/boneres.2016.4 (2016).

13. Lee, W. C., Ji, X., Nissim, I. & Long, F. Malic Enzyme Couples Mitochondria with Aerobic Glycolysis in Osteoblasts. Cell Rep 32, 108108, doi:10.1016/j.celrep.2020.108108 (2020).

14. Guntur, A. R., Le, P. T., Farber, C. R. & Rosen, C. J. Bioenergetics during calvarial osteoblast differentiation reflect strain differences in bone mass. Endocrinology 155, 1589–1595, doi:10.1210/en.2013-1974 (2014).

15. Bonds, D. E. et al. Risk of fracture in women with type 2 diabetes: the Women’s Health Initiative Observational Study. J Clin Endocrinol Metab 91, 3404–3410, doi:10.1210/jc.2006-0614 (2006).

16. Ma, L. et al. Association between bone mineral density and type 2 diabetes mellitus: a meta-analysis of observational studies. Eur J Epidemiol 27, 319–332, doi:10.1007/s10654-012-9674-x (2012).

17. Sellmeyer, D. E. et al. Skeletal Metabolism, Fracture Risk, and Fracture Outcomes in Type 1 and Type 2 Diabetes. Diabetes 65, 1757–1766, doi:10.2337/db16-0063 (2016).

18. Farr, J. N. & Khosla, S. Determinants of bone strength and quality in diabetes mellitus in humans. Bone 82, 28–34, doi:10.1016/j.bone.2015.07.027 (2016).

19. Li, Z. et al. The relationship between estimated glucose disposal rate and bone turnover markers in type 2 diabetes mellitus. Endocrine 77, 242–251, doi:10.1007/s12020-022-03090-z (2022).

20. Moseley, K. F., Du, Z., Sacher, S. E., Ferguson, V. L. & Donnelly, E. Advanced glycation endproducts and bone quality: practical implications for people with type 2 diabetes. Curr Opin Endocrinol Diabetes Obes 28, 360–370, doi:10.1097/MED.0000000000000641 (2021).

21. Nadeau, K. J. et al. Youth-Onset Type 2 Diabetes Consensus Report: Current Status, Challenges, and Priorities. Diabetes Care 39, 1635–1642, doi:10.2337/dc16-1066 (2016).

22. Kindler, J. M. et al. Bone Mass and Density in Youth With Type 2 Diabetes, Obesity, and Healthy Weight. Diabetes Care 43, 2544–2552, doi:10.2337/dc19-2164 (2020).

23. Song, D. et al. Inducible expression of Wnt7b promotes bone formation in aged mice and enhances fracture healing. Bone Res 8, 4, doi:10.1038/s41413-019-0081-8 (2020).

24. Pereira, R. O. et al. Inducible overexpression of GLUT1 prevents mitochondrial dysfunction and attenuates structural remodeling in pressure overload but does not prevent left ventricular dysfunction. J Am Heart Assoc 2, e000301, doi:10.1161/JAHA.113.000301 (2013).

25. Kim, W. Y. et al. Failure to prolyl hydroxylate hypoxia-inducible factor alpha phenocopies VHL inactivation in vivo. EMBO J 25, 4650–4662, doi:10.1038/sj.emboj.7601300 (2006).

26. Perl, A. K., Wert, S. E., Nagy, A., Lobe, C. G. & Whitsett, J. A. Early restriction of peripheral and proximal cell lineages during formation of the lung. Proc Natl Acad Sci U S A 99, 10482–10487 (2002).

27. Melamud, E., Vastag, L. & Rabinowitz, J. D. Metabolomic analysis and visualization engine for LC-MS data. Anal Chem 82, 9818–9826, doi:10.1021/ac1021166 (2010).

28. Stuart, T. et al. Comprehensive Integration of Single-Cell Data. Cell 177, 1888–1902 e1821, doi:10.1016/j.cell.2019.05.031 (2019).

29. Baccin, C. et al. Combined single-cell and spatial transcriptomics reveal the molecular, cellular and spatial bone marrow niche organization. Nat Cell Biol 22, 38–48, doi:10.1038/s41556-019-0439-6 (2020).

30. Morikawa, S. et al. Prospective identification, isolation, and systemic transplantation of multipotent mesenchymal stem cells in murine bone marrow. J Exp Med 206, 2483–2496, doi:10.1084/jem.20091046 (2009).

31. Baryawno, N. et al. A Cellular Taxonomy of the Bone Marrow Stroma in Homeostasis and Leukemia. Cell 177, 1915–1932 e1916, doi:10.1016/j.cell.2019.04.040 (2019).

32. Moonira, T. et al. Metformin lowers glucose 6-phosphate in hepatocytes by activation of glycolysis downstream of glucose phosphorylation. J Biol Chem 295, 3330–3346, doi:10.1074/jbc.RA120.012533 (2020).

33. Valero, T. Mitochondrial biogenesis: pharmacological approaches. Curr Pharm Des 20, 5507–5509, doi:10.2174/138161282035140911142118 (2014).

34. Wang, Y. et al. Metformin Improves Mitochondrial Respiratory Activity through Activation of AMPK. Cell Rep 29, 1511–1523 e1515, doi:10.1016/j.celrep.2019.09.070 (2019).

35. Zhu, X. et al. Metformin improves cognition of aged mice by promoting cerebral angiogenesis and neurogenesis. Aging (Albany NY) 12, 17845–17862, doi:10.18632/aging.103693 (2020).

36. Wei, J. et al. Glucose Uptake and Runx2 Synergize to Orchestrate Osteoblast Differentiation and Bone Formation. Cell 161, 1576–1591, doi:10.1016/j.cell.2015.05.029 (2015).

37. De Bock, K. et al. Role of PFKFB3-driven glycolysis in vessel sprouting. Cell 154, 651–663, doi:10.1016/j.cell.2013.06.037 (2013).

38. Napoli, N. et al. Mechanisms of diabetes mellitus-induced bone fragility. Nat Rev Endocrinol 13, 208–219, doi:10.1038/nrendo.2016.153 (2017).

39. Brownlee, M. Biochemistry and molecular cell biology of diabetic complications. Nature 414, 813–820, doi:10.1038/414813a (2001).

